# Multiple evolutionary origins reflect the importance of sialic acid transporters in the colonisation potential of bacterial pathogens and commensals

**DOI:** 10.1101/2021.03.01.433349

**Authors:** Emmanuele Severi, Michelle Rudden, Andrew Bell, Tracy Palmer, Nathalie Juge, Gavin H Thomas

## Abstract

Located at the tip of cell surface glycoconjugates, sialic acids are at the forefront of host-microbe interactions and, being easily liberated by sialidase enzymes, are used as metabolites by numerous bacteria, particularly by pathogens and commensals living on or near diverse mucosal surfaces. These bacteria rely on specific transporters for the acquisition of host-derived sialic acids. Here, we present the first comprehensive genomic and phylogenetic analysis of bacterial sialic acid transporters, leading to the identification of multiple new families and subfamilies. Our phylogenetic analysis suggests that sialic acid-specific transport has evolved independently at least 8 times during the evolution of bacteria, from within 4 of the major families/superfamilies of bacterial transporters, and we propose a robust classification scheme to bring together a myriad of different nomenclatures that exist to date. The new transporters discovered occur in diverse bacteria including Spirochaetes, Bacteroidetes, Planctomycetes, and Verrucomicrobia, many of which are species that have not been previously recognised to have sialometabolic capacities. Two subfamilies of transporters stand out in being fused to the sialic acid mutarotase enzyme, NanM, and these transporter fusions are enriched in bacteria present in gut microbial communities. We also provide evidence for a possible function of a sialic acid transporter component in chemotaxis that is independent of transport. Our analysis supports the increasing experimental evidence that competition for host-derived sialic acid is a key phenotype for successful colonisation of complex mucosal microbiomes, such that a strong evolutionary selection has occurred for the emergence of sialic acid specificity within existing transporter architectures.

## INTRODUCTION

“Sialic acid” is a generic term covering a family of over 50 related sugar acids that are ubiquitous among vertebrates, where they occur as terminal sugars of cell surface-exposed glycoconjugates in several tissues [1–4]. The most commonly studied sialic acid, *N*-acetyl-neuraminic acid (Neu5Ac, Fig. 1), is the only form synthesised by humans, whereas other animals can also make C5-group variants such as *N*-glycolyl-neuraminic (Neu5Gc) and 3-keto-3-deoxy-D-glycero-D-galactonononic (KDN) acids [2, 3, 5]. Various *O*-substitutions at different hydroxyl moieties expand the diversity of sialic acid in nature [2]. The location of structurally diverse sialic acids at the cellular surface underpins the wide variety of biological roles this molecule plays in the host. These often rely on physical interactions between sialylated glycoproteins and various partners, and they include embryonic development and regulation of the immune system, where sialic acid contributes to the recognition of the “self” [1, 3].

**Figure 1.**
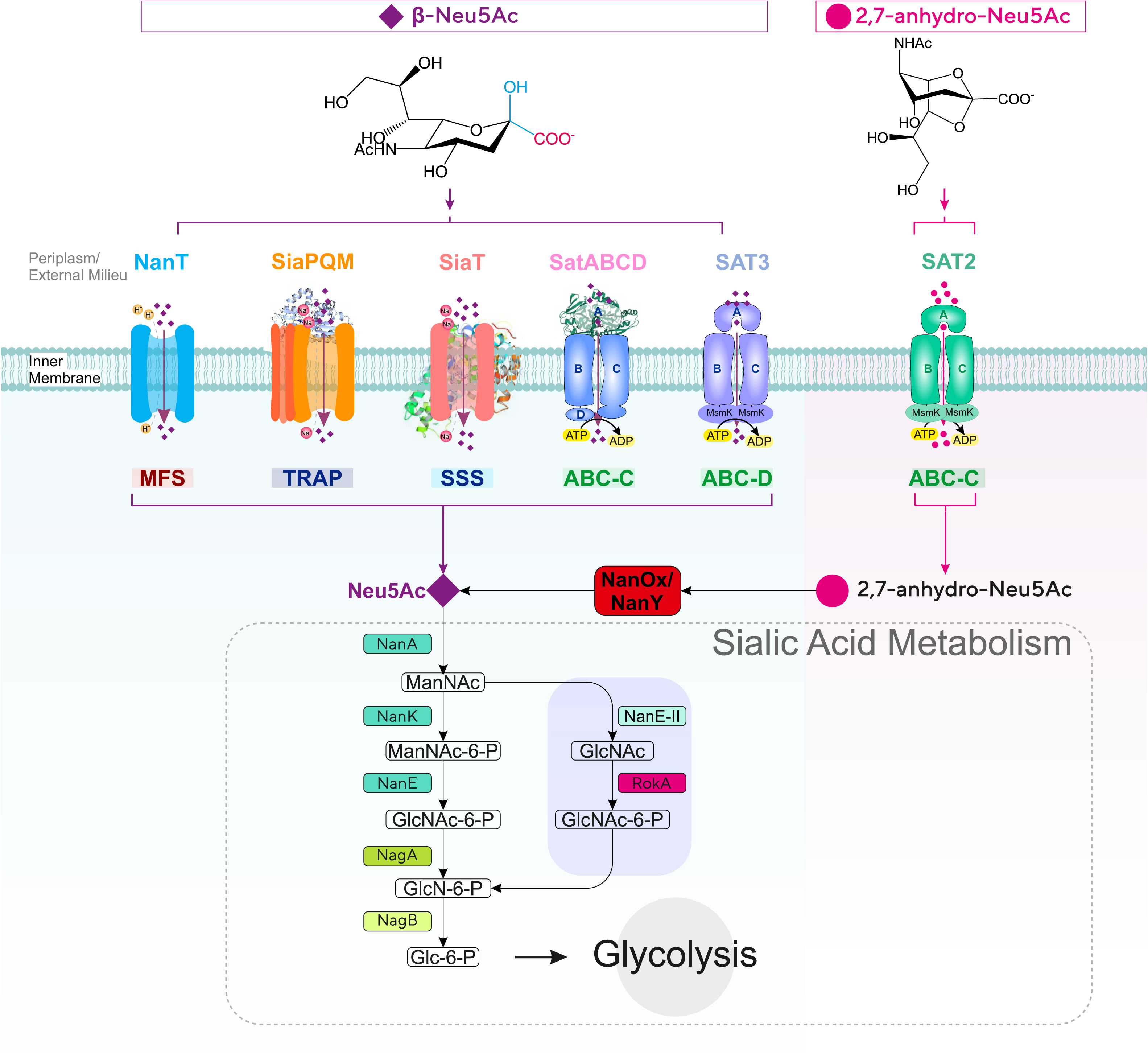
Diversity of sialic acid transport and catabolism in bacteria. Sialic acid-utilising bacteria have evolved multiple types of transporters functioning at the inner (cytoplasmic) membrane from 4 major (super)families (MFS, TRAP, ABC, and SSS) differing by mode of energisation and subunit composition. Once inside the cell, sialic acid is metabolised to GlcNAc-6-P *via* one of two characterised pathways to then enter central metabolism by the action of the GlcNAc-catabolic enzymes NagA (GlcNAc-6-P deacetylase) and NagB (GlcN-6-P deaminase). In the *E. coli* paradigm [27] Neu5Ac is converted by the sequential action of three dedicated enzymes, namely NanA (Neu5Ac lyase), NanK (ManNAc kinase), and NanE (ManNAc-6-P epimerase); in the Bacteroidetes paradigm [30, 31] NanA and the alternative enzyme NanE-II (GlcNAc epimerase) feed substrate to the glucokinase RokA. Bacteria such as *R. gnavus* that take up 2,7-anhydro-Neu5Ac use the cytoplasmic oxidoreductase NanOx (NanY in *E. coli*) to convert the substrate to Neu5Ac, which then enters a canonical Nan pathway. ManNAc: N-acetyl-mannosamine; GlcNAc: N-acetyl-glucosamine; GlcN: glucosamine; Glc: glucose. MsmK: multitasking ATP-ase protein serving multiple sugar ABC transporters including sialic acid transporters [44]. SusCD: outer membrane protein complex for glycan acquisition made of a TonB-dependent transporter (SusC) and an extracytpolasmic lipoprotein (SusD). NanOU is an experimentally confirmed sialic acid-specific SusCD-family complex [83]. MFS: major facilitator superfamily. ABC: ATP-binding cassette. TRAP: tripartite ATP-independent periplasmic (transporters). SSS: sodium-solute symporters.

Being surface located also means that sialic acid comes into direct contact with microorganisms at the host mucosal surface. While human viruses such as influenza and MERS coronavirus are most (in)famous for their ability to target sialylated receptors to cause disease [6, 7], bacteria can also utilise sialic acid as a mediator of interactions with the host [4, 8–10]. Both partnership-establishing bacteria (i.e., symbionts or commensals) and pathogens can release and metabolise host sialic acid, which can then be incorporated into surface structures such as capsule and LPS that may help evade the host’s immune system through so-called “molecular mimicry”, i.e., by hijacking the role of sialic acid in the host’s recognition of the self. Further, sialic acid can be used as a metabolic substrate to sustain growth and enhance the ability of bacterial species to establish themselves in target niches, in health or disease [4, 8–10].

While a small minority of sialic acid-utilising microbes can synthesise Neu5Ac *de novo* [11], all others rely on host-derived sialic acid, whether for growth or cell surface sialylation [4, 8, 9], acquired through dedicated sialic acid transporters [8, 10, 12]. Although not ubiquitous among prokaryotes, sialic acid transport is a widespread trait across all types of sialic acid-utilising bacteria that predominantly inhabit mucosal surfaces [4, 5, 8, 13], where it plays a critical role in virulence and host colonisation [9, 14–19]. The role of sialic acid transporters in host-microbe interactions has been the target of extensive research including from ourselves [4, 8, 10, 12, 13, 20], and today six types of bacterial sialic acid transporters have been characterised experimentally (Fig. 1) [12, 18]. These uptake systems differ by a number of features including mode of energisation, subunit composition, and substrate specificity (Fig. 1 and [12]), indicating that sialic acid transport has evolved independently multiple times and that there is selective pressure for the acquisition of this trait in numerous, taxonomically diverse prokaryotes [12].

The first five types of transporters to have been studied all include systems specific for Neu5Ac (Fig. 1), with some able to take up Neu5Gc or KDN [9, 20–24]. The MFS (Major Facilitator Superfamily) transporter, NanT was the first sialic acid transporter reported during ground-breaking work by the Vimr group on Neu5Ac catabolism in *E. coli* [25, 26], which also elucidated the canonical prokaryotic metabolism pathway for sialic acid comprising Nan and Nag enzymes [27] (Fig. 1). As an MFS protein, NanT is a secondary transporter that uses the proton gradient to drive concentrative uptake of Neu5Ac, and has been studied both *in vivo* and *in vitro* in some detail [21, 25, 28, 29]. The MFS transporter from the Bacteroidetes, *Bacteroides fragilis* and *Tannerella forsythia*, also called NanT, is normally considered as being of the same family as the Enterobacterial NanT group, despite notable differences between it and *E. coli* NanT [12, 30, 31]. The TRAP (tripartite ATP-independent periplasmic) transporter SiaPQM, first characterised in *Haemophilus influenzae* [14, 32] and also well-studied in *Vibrio* spp. [28, 33, 34], is among the best-studied sialic acid transporters to date, with a wealth of *in vivo* and *in vitro* data based on the use of native and heterologous hosts as well as reconstituted purified systems [14, 28, 35, 36]. SiaPQM too is a secondary transporter, being energised by a sodium rather than a proton gradient [29, 36], but it also depends on the solute-binding protein (SBP), SiaP, for function [14, 35]. While use of an SBP is a feature most commonly associated with primary (e.g. ATP-dependent) transporters [35], SiaP here functions together with two transmembrane (TMH) components that bear no relationships with those of primary systems (the 4 TMH SiaQ and the 12 TMH SiaM, fused in *H. influenzae*) [37]. Discovered the same year as SiaPQM, SatABCD is another SBP-dependent system, but from the ABC (ATP-binding cassette) transporter superfamily, i.e., a primary transporter that uses ATP binding and hydrolysis to energise uptake. The best characterised SatABCD system to date is that of *Haemophilus ducreyi* (HdSatABCD) [38], but genetic studies have been carried out in Actinobacteria too [39, 40]. As also reported for SiaP [35, 41–43], crystallographic and mutational studies identified key residues for Neu5Ac-binding and selectivity by the SBP component, SatA [23]. Not to be confused with the *H. ducreyi* transporter is a different ABC system discovered in *Streptococcus pneumoniae* [16], normally referred to as SatABC too, but also as SAT3 [13]. To date information on this transporter derives solely from genetic and transcriptional studies [16, 44–46] and, despite the distinction from HdSatABCD dating back to 2009 [13], this system remains to be functionally characterised. The fifth group of Neu5Ac-specific transporters include proteins of the SSS (sodium-solute symporter) family of secondary transporters, which are generally referred to as “SiaT” [12, 22, 47]. First discovered about 10 years ago as a diverse and widespread group of transporters [13, 48], several SiaT transporters from Gram-negative and Gram-positive bacteria have now been functionally characterised *in vivo* and *in vitro* [4, 22, 47, 48], and a high-resolution crystal structure is available for the *Proteus mirabilis* orthologue with bound Neu5Ac [47]. Complementation, mutagenesis, and biochemical studies using reconstituted systems confirmed the dependence of these proteins on sodium for function, identified residues involved in substrate-binding and transport of coupled Na^+^ ions, and provided insight into their substrate specificity [21, 22, 47, 48].

The sixth and final group of characterised sialic acid transporters has introduced an entirely novel substrate specificity to the field. Early gene cluster analyses had predicted that ABC uptake systems of the “SAT2” type were sialic acid transporters distinct from the above HdSatABCD (“SAT”) and SAT3 ABC systems [13], but SAT2 transporters remained uncharacterised until recent work [18] studying strains of the gut symbiont *Ruminococcus gnavus*, which have the capacity to produce a unique anhydro form of sialic acid (2,7-anhydro-Neu5Ac, Fig. 1) from Neu5Ac-terminated glycoconjugates *via* the action of an intramolecular *trans*-sialidase (IT-sialidase) [49, 50]. Working with mutant strains as well as with the purified SBP component of SAT2, Bell *et al.* [18] demonstrated that the *R. gnavus* SBP (*Rg*SBP) was specific for 2,7-anhydro-Neu5Ac (Fig. 1). The study established that the IT-sialidase, the SAT2 transporter, and the newly discovered oxidoreductase, NanOx, which converts 2,7-anhydro-Neu5Ac back to Neu5Ac before being metabolised further (Fig. 1; [18, 51]), cooperate to channel an “exclusive” form of sialic acid in to an otherwise canonical Nan catabolic pathway [18] (Fig. 1). This is a compelling instance where prokaryotes have innovated on sialic acid transport and used it to their advantage in their target niche [18, 49]. To date, this is the sole example for this group of ABC sialic acid transporters [16]. However, the characterisation of the orthologous oxidoreductases NanOx from *R. gnavus* and NanY (formerly YjhC) from *E. coli* [51, 52], has provided evidence for at least three other potential anhydro-sialic acid transporters. Functional complementation of *E. coli* mutants has demonstrated a role in anhydro-sialic acid acquisition for one of these, namely the NanT-like *E. coli* MFS transporter, NanX (formerly YjhB) [52, 53].

The diversity of bacterial sialic acid transporters has raised various questions regarding structural-functional features, mechanism of transport, and exact physiological roles [12, 54]. However, sialic acid transport has seldom been approached from a phylogenetic/evolutionary perspective, in stark contrast with the rich ensemble of phylogenetic studies on the Nan catabolic enzymes, namely NanA, NanE, and NanK (Fig. 1; [9, 13, 30, 55]). As these studies have revealed the existence of diverse clades of NanA, NanE, and NanK orthologues often combining into mosaic clusters at gene level, equivalent analyses are largely missing for the accompanying transporters, with the sole exception of studies on SiaP [33]. There is a need to update the broad classification of sialic acid transporters put forward by the Boyd group in 2009 [13] taking into account the distribution of individual uptake systems across bacteria [14, 20, 47, 48].

Here, we carried out gene cluster and phylogenetic analyses of all known sialic acid transporters across the bacteria. Using phylogeny as the basis for classification, we first demonstrated that all described sialic acid transporters can be classified into eight distinct “families”, with validation of the six historically established types (Fig. 1), and with the identification of two new families of phylogenetically distinct MFS transporters. Within the SSS and MFS families we discovered a novel form of sialic acid transporter, which consists of a fusion with an enzyme involved in its processing and is widespread among the Bacteroidetes and Planctomycetes/Verrucomicrobia. Overall, the study provides significant new insights into the evolution and function of sialic acid transporters, as well as revealing other novel aspects, including a potential novel biological role in chemotaxis for a sialic acid transporter component.

## RESULTS

### Phylogenetic analyses identify eight different evolutionary origins for bacterial sialic acid transporters

In light of our recent discovery of two new 2,7-anhydro-sialic acid transporters [18, 51], we undertook the first systematic bioinformatic analysis of sialic acid transporters across bacteria. The approach used, described in (13), is based on the genetic association of a transporter gene with a complete *nan* catabolic pathway (see Methods) as the primary factor for inclusion. Examples of characterised bacterial *nan* clusters are shown in Fig. 2, with a more detailed results of the analysis presented in Supplementary Figures S2-S12. This bioinformatics approach revealed expanded groups for all known types of sialic acid transporters as well as additional novel groups linked to known *nan* genes. Within both the MFS superfamily and the large SSS family of transporters there were potentially multiple sub-families of sialic acid transporters, and their mono- or polyphyletic origin was investigated using a phylogenetic approach (Fig. 3). For the MFS proteins in Pfam family MFS_1, two different evolutionary origins of sialic acid transporters were clearly identified (Fig. 3A), and a third origin was discovered uniquely in the MFS_2 family, which encompassed the GPH (Glycoside-Pentoside-Hexuronide:cation symporter) family [56, 57] (Fig. 3B). This contrasts with the SSS transporters, where 3 potentially different families appear to form a clear monophyletic group, suggesting that sialic acid specificity emerged once and then these proteins have diversified into related clades (Fig. 3C). For the SBP-dependent transporters, the phylogenetic analysis targeted the SBP component, which defines the substrate-specificity, and based on this principle we found a single origin for the TRAP transporters, as previously suggested [33] (Fig. 3D). As for the ABC transporters, which fall into two Pfam families, there is evidence of two independent origins for Pfam SBP_bac_1 (Cluster C SBPs) (Fig. 3E), while a single origin was identified for the SBP_bac_5 (Cluster D SBPs) (Fig. 3F). From this analysis, we can conclude that sialic acid transport specificity has evolved at least 8 times during the evolution of bacteria, and to aid in the classification of these diverse transporters we have named these using a sialic acid transporter (ST) family nomenclature, from ST1-8 (Table 1).

**Figure 2.**
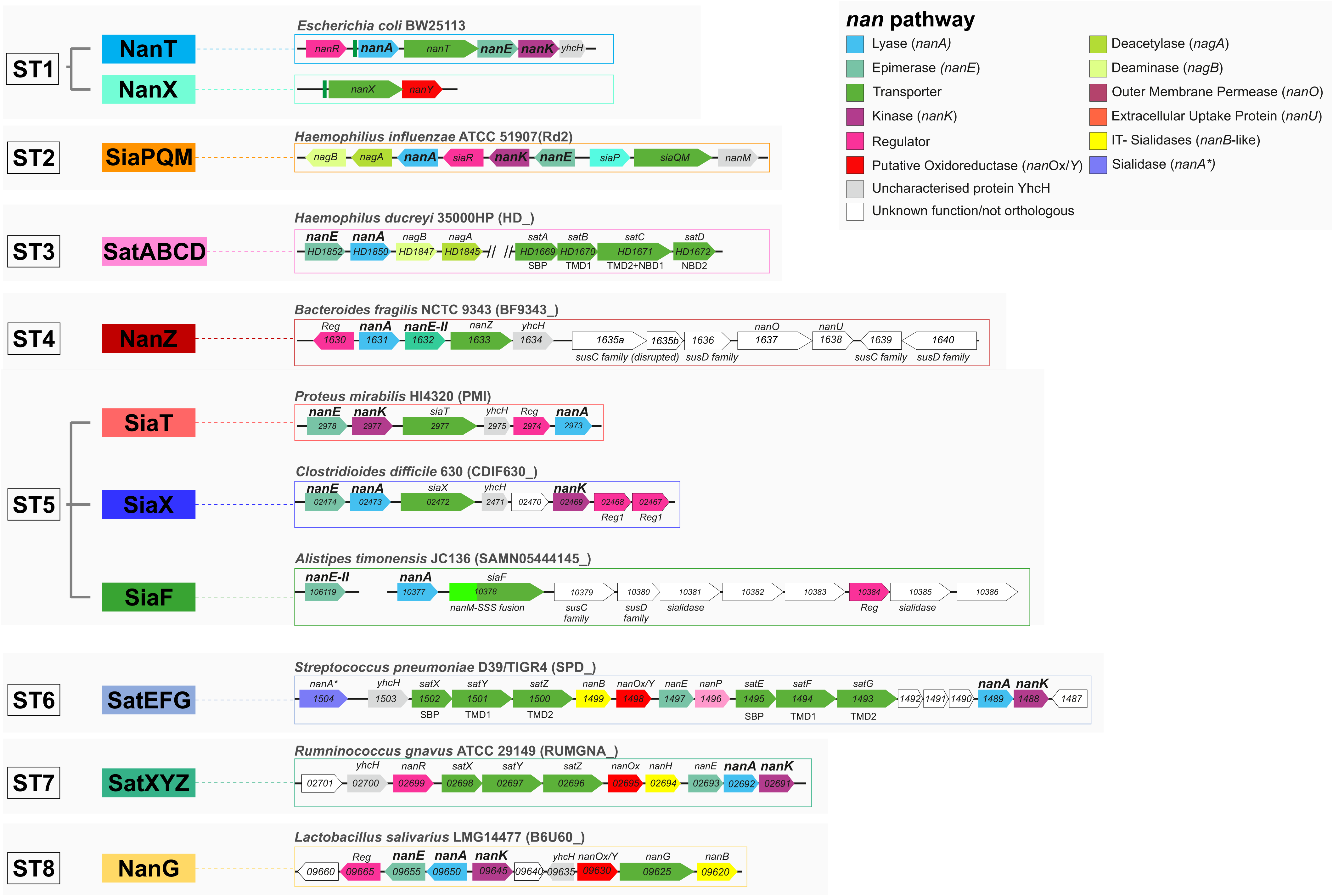
Structure of *nan* clusters for archetypal transporter families. *nan* clusters are shown for each of the eight newly classified sialic acid transporter families using archetypical organisms as reference. Locus prefixes are denoted in brackets next to reference organism, gene names are displayed within gene tags. *nanOp* operators highlighted upstream of *nanA* and *nanX* in the *E. coli* ST1 loci emphasise the occurrence of a single NanR regulon in *E. coli* [53]. In TIGR4, the ST6 locus bears minor differences in the form of pseudogenes and insertions [53]. Note that, as Post et al [38] before us, we could not find orthologues of *nanK* in *H. ducreyi* 35000HP, thus ManNAc kinase functions in this organism remain unidentified.

**Figure 3.**
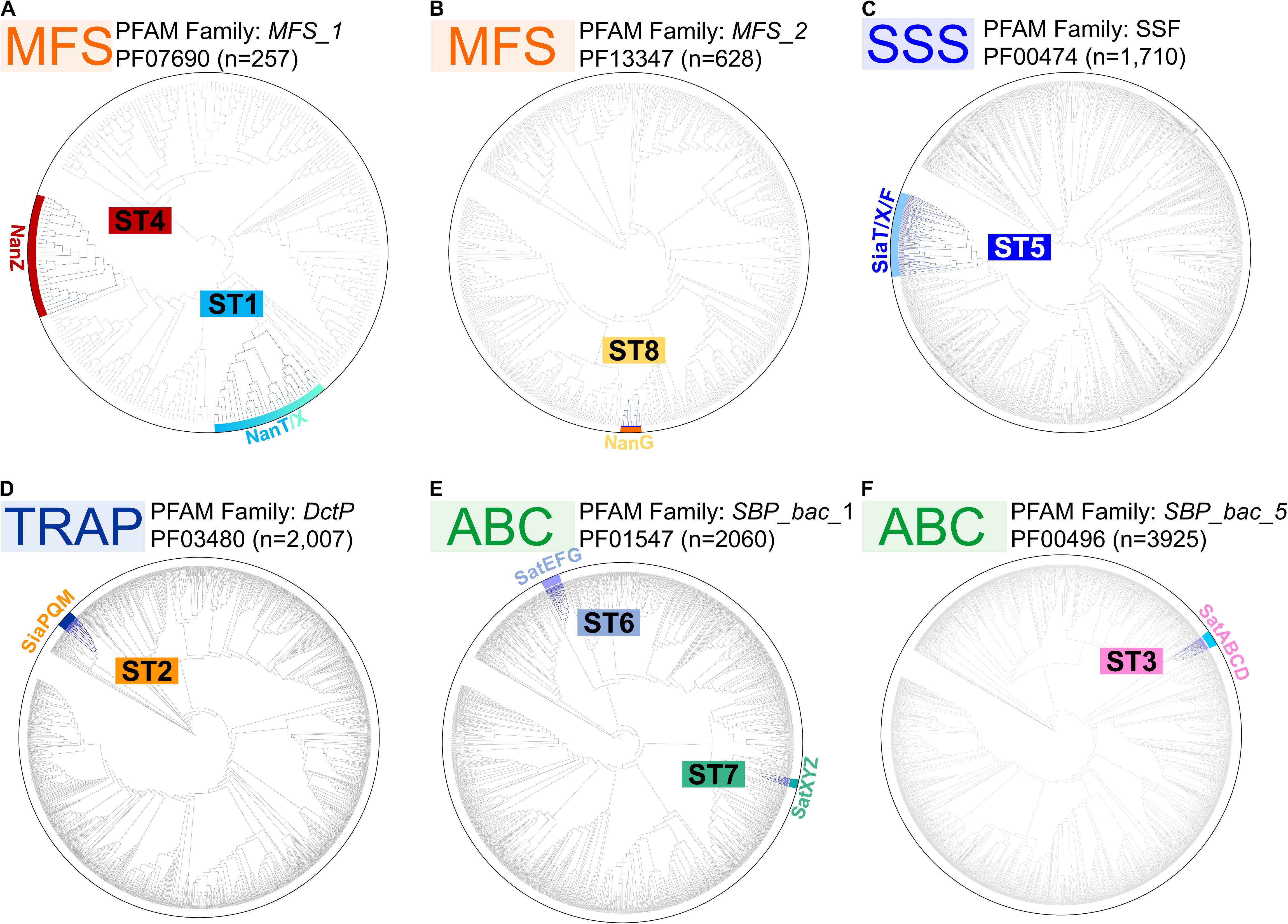
Phylogenetic classification of sialic acid transporters in bacteria. Global phylogenetic analysis of sialic acid transporters within expanded Pfam families. Clades highlighted in blue are sialic acid transporter sequences that reside in verified *nan* clusters. Among the diverse MFS_1 and ABC (*SBP_bac_1 -* Cluster C SBPs) families we observe independent evolutionary origins. In contrast, in the SSS, TRAP, and ABC (*SBP_bac_5* - Cluster D SBPs) families appear each to have only single ST origins.

**Table 1.**
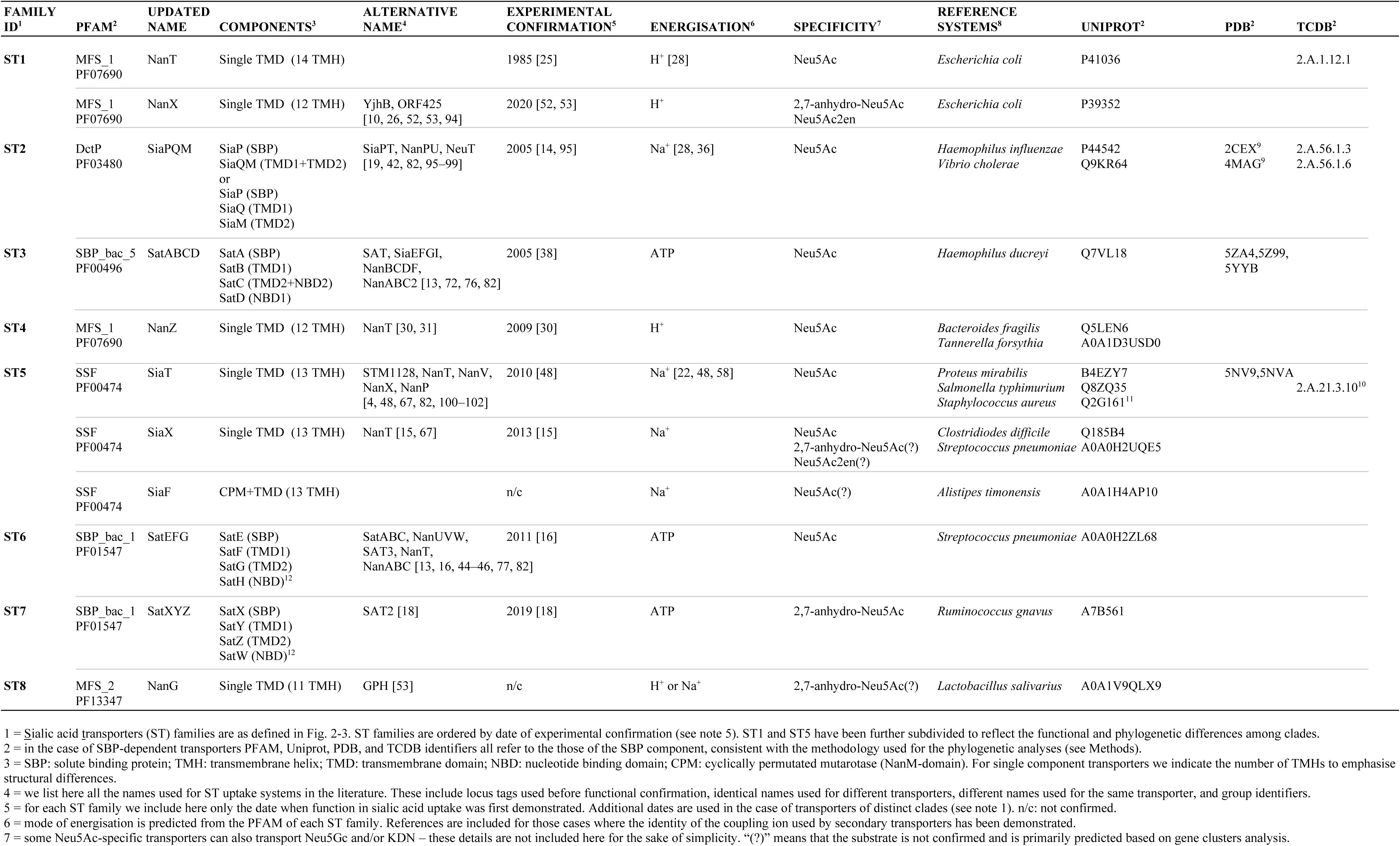
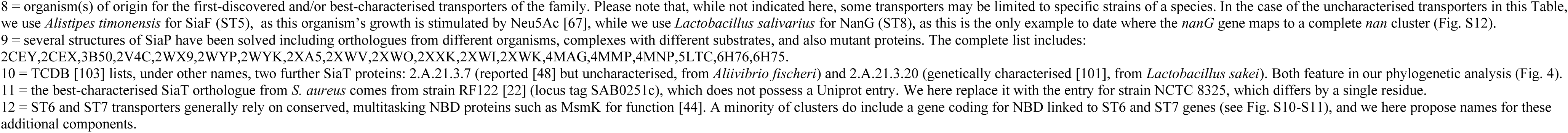
Classification and characteristics of bacterial sialic acid transporters

| FAMILY ID<sup>1</sup> | PFAM<sup>2</sup> | UPDATED NAME | COMPONENTS<sup>3</sup> | ALTERNATIVE NAME<sup>4</sup> | EXPERIMENTAL CONFIRMATION<sup>5</sup> | ENERGISATION<sup>6</sup> | SPECIFICITY<sup>7</sup> | REFERENCE SYSTEMS<sup>8</sup> | UNIPROT<sup>2</sup> | PDB<sup>2</sup> | TCDB<sup>2</sup> |
| --- | --- | --- | --- | --- | --- | --- | --- | --- | --- | --- | --- |
| ST1 | MFS_1<br>PF07690 | NanT | Single TMD (14 TMH) |  | 1985 [25] | H<sup>+</sup> [28] | Neu5Ac | *Escherichia coli* | P41036 |  | 2.A.1.12.1 |
|  | MFS_1<br>PF07690 | NanX | Single TMD (12 TMH) | YjhB, ORF425<br>[10, 26, 52, 53, 94] | 2020 [52, 53] | H<sup>+</sup> | 2,7-anhydro-Neu5Ac<br>Neu5Ac2en | *Escherichia coli* | P39352 |  |  |
| ST2 | DctP<br>PF03480 | SiaPQM | SiaP (SBP)<br>SiaQM (TMD1+TMD2)<br>or<br>SiaP (SBP)<br>SiaQ (TMD1)<br>SiaM (TMD2) | SiaPT, NanPU, NeuT<br>[19, 42, 82, 95–99] | 2005 [14, 95] | Na<sup>+</sup> [28, 36] | Neu5Ac | *Haemophilus influenzae*<br>*Vibrio cholerae* | P44542<br>Q9KR64 | 2CEX<sup>9</sup><br>4MAG<sup>9</sup> | 2.A.56.1.3<br>2.A.56.1.6 |
| ST3 | SBP_bac_5<br>PF00496 | SatABCD | SatA (SBP)<br>SatB (TMD1)<br>SatC (TMD2+NBD2)<br>SatD (NBD1) | SAT, SiaEFGI,<br>NanBCDF,<br>NanABC2 [13, 72, 76, 82] | 2005 [38] | ATP | Neu5Ac | *Haemophilus ducreyi* | Q7VL18 | 5ZA4,5Z99,<br>5YYB |  |
| ST4 | MFS_1<br>PF07690 | NanZ | Single TMD (12 TMH) | NanT [30, 31] | 2009 [30] | H<sup>+</sup> | Neu5Ac | *Bacteroides fragilis*<br>*Tannerella forsythia* | Q5LEN6<br>A0A1D3USD0 |  |  |
| ST5 | SSF<br>PF00474 | SiaT | Single TMD (13 TMH) | STM1128, NanT, NanV,<br>NanX, NanP<br>[4, 48, 67, 82, 100–102] | 2010 [48] | Na<sup>+</sup> [22, 48, 58] | Neu5Ac | *Proteus mirabilis*<br>*Salmonella typhimurium*<br>*Staphylococcus aureus* | B4EZY7<br>Q8ZQ35<br>Q2G161<sup>11</sup> | 5NV9,5NVA | 2.A.21.3.10<sup>10</sup> |
|  | SSF<br>PF00474 | SiaX | Single TMD (13 TMH) | NanT [15, 67] | 2013 [15] | Na<sup>+</sup> | Neu5Ac<br>2,7-anhydro-Neu5Ac(?)<br>Neu5Ac2en(?) | *Clostridiodes difficile*<br>*Streptococcus pneumoniae* | Q185B4<br>A0A0H2UQE5 |  |  |
|  | SSF<br>PF00474 | SiaF | CPM+TMD (13 TMH) |  | n/c | Na<sup>+</sup> | Neu5Ac(?) | *Alistipes timonensis* | A0A1H4AP10 |  |  |
| ST6 | SBP_bac_1<br>PF01547 | SatEFG | SatE (SBP)<br>SatF (TMD1)<br>SatG (TMD2)<br>SatH (NBD)<sup>12</sup> | SatABC, NanUVW,<br>SAT3, NanT,<br>NanABC [13, 16, 44–46, 77, 82] | 2011 [16] | ATP | Neu5Ac | *Streptococcus pneumoniae* | A0A0H2ZL68 |  |  |
| ST7 | SBP_bac_1<br>PF01547 | SatXYZ | SatX (SBP)<br>SatY (TMD1)<br>SatZ (TMD2)<br>SatW (NBD)<sup>12</sup> | SAT2 [18] | 2019 [18] | ATP | 2,7-anhydro-Neu5Ac | *Ruminococcus gnavus* | A7B561 |  |  |
| ST8 | MFS_2<br>PF13347 | NanG | Single TMD (11 TMH) | GPH [53] | n/c | H<sup>+</sup> or Na<sup>+</sup> | 2,7-anhydro-Neu5Ac(?) | *Lactobacillus salivarius* | A0A1V9QLX9 |  |  |

= in the case of SBP-dependent transporters PFAM, Uniprot, PDB, and TCDB identifiers all refer to the those of the SBP component, consistent with the methodology used for the phylogenetic analyses (see Methods).
= SBP: solute binding protein; TMH: transmembrane helix; TMD: transmembrane domain; NBD: nucleotide binding domain; CPM: cyclically permuted mutarotase (NanM-domain). For single component transporters we indicate the number of TMHs to emphasise structural differences.
= we list here all the names used for ST uptake systems in the literature. These include locus tags used before functional confirmation, identical names used for different transporters, different names used for the same transporter, and group identifiers.
= for each ST family we include here only the date when function in sialic acid uptake was first demonstrated. Additional dates are used in the case of transporters of distinct clades (see note 1). n/c: not confirmed.
= mode of energisation is predicted from the PFAM of each ST family. References are included for those cases where the identity of the coupling ion used by secondary transporters has been demonstrated.
= some Neu5Ac-specific transporters can also transport Neu5Gc and/or KDN – these details are not included here for the sake of simplicity. “(?)” means that the substrate is not confirmed and is primarily predicted based on gene clusters analysis.
= several structures of SiaP have been solved including orthologues from different organisms, complexes with different substrates, and also mutant proteins. The complete list includes: 2CEY, 2CEX, 3B50, 2V4C, 2WX9, 2WYP, 2WYK, 2XA5, 2XWV, 2XWO, 2XXK, 2XWI, 2XWK, 4MAG, 4MMP, 4MNP, 5LTC, 6H76, 6H75.
= the best-characterised SiaT orthologue from *S. aureus* comes from strain RF122 [22] (locus tag SAB0251c), which does not possess a Uniprot entry. We here replace it with the entry for strain NCTC 8325, which differs by a single residue.

### The ST5 (SSS) sialic acid transporters cluster into distinct clades and include candidate anhydro-sialic acid transporters

Although the SSS transporters identified in our *in-silico* analyses appear to have a single evolutionary origin (ST5) (Fig. 3C), we noticed that they are the most widespread in bacteria, and we could see distinct clades within them (Fig. 4). Previously characterised ST5 sialic acid transporters [15, 22, 48, 58] map to two of three major clades. The “SiaT” clade contains the SiaT proteins from Enterobacteria and *S. aureus*, while the second clade contains the SSS transporter from *Clostridioides difficile*, which, having previously been called NanT (Table 1), we propose instead to name SiaX to both stress its ST5 nature and emphasise the distinction among ST5 clades (Fig. 4). The SiaT group has the broadest phylogenetic distribution, while the SiaX group is limited to Firmicutes (Fig. 4). A third major clade, to which we refer as “SiaF” (Fig. 4), contains exclusively newly predicted sialic acid transporters (see later section). Of note within the SiaT clade are a novel sub-clade occurring in some *Brachyspira* spp. (see next section) and a smaller branch featuring transporters encoded within *nan* clusters of *Mycoplasma* spp. [13, 59].

**Figure 4.**
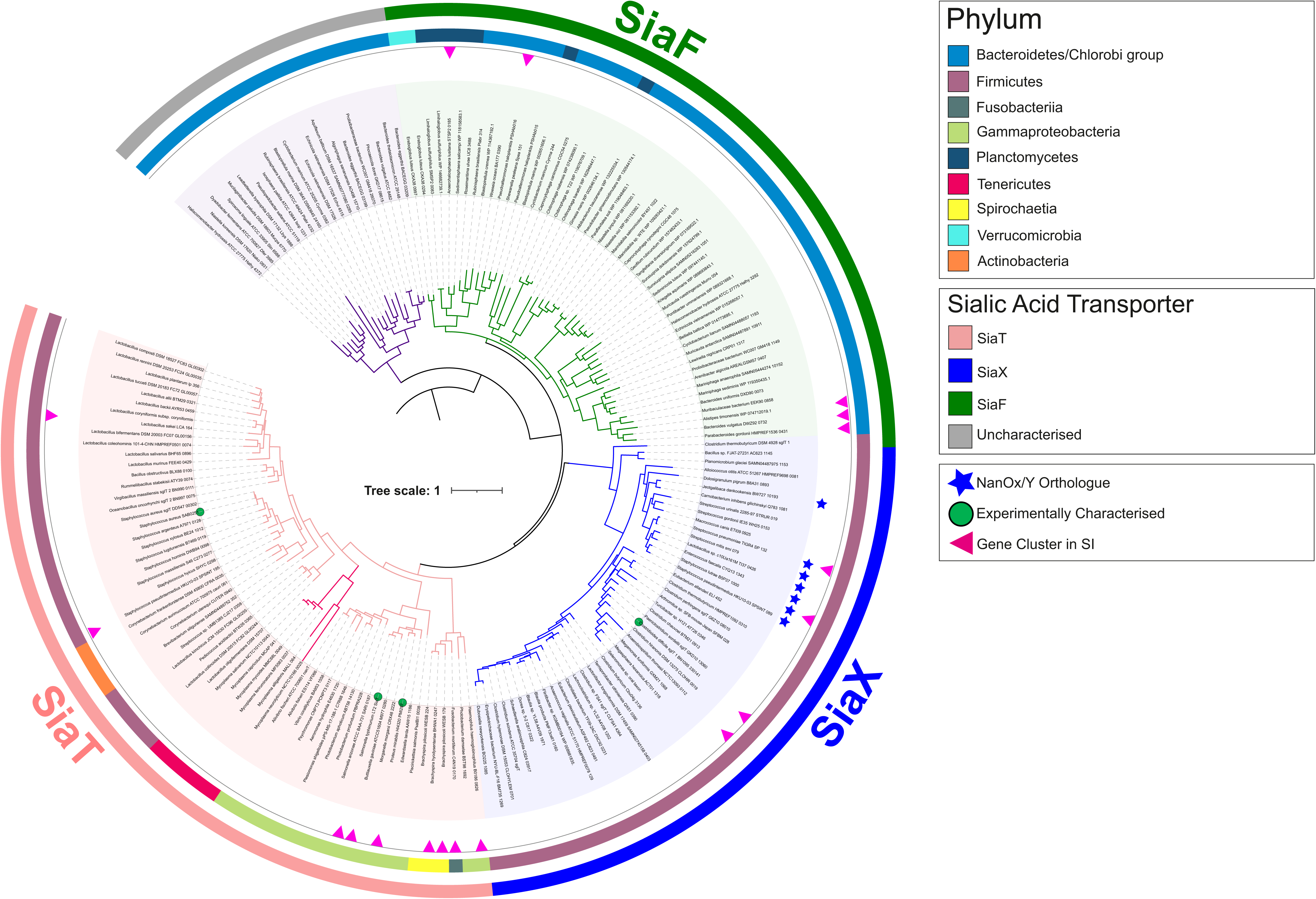
Phylogenetic distribution of ST5 (SSS) sialic acid transporters in bacteria. Phylogeny of ST5 (SSS) sialic acid transporters at phylum level. Coloured branches represent three major clades, the historical SiaT clade (red), the SiaX clade (blue) including both Neu5Ac and putative anhydro-Neu5Ac transporters (asterisk), and the clade of SiaF proteins (green) representing a novel fusion between sialic acid transporter and mutarotase. A fourth group of uncharacterised transporters (grey) including one from *Bacteroides thetaiotaomicron* is addressed in the Discussion. Experimentally characterised transporters are highlighted on the tree with a red circle. SiaT is distributed widely across several bacterial phyla, whereas SiaX is restricted to the Firmicutes, and SiaF occurs near-exclusively across *Bacteroides* and *Planctomycetes/Verrucomicrobia*. Maximum likelihood tree was inferred from SSS transporter proteins(n=354) residing within a *nan* cluster containing at least one sialocatabolic *nan* gene.

Notable with the SiaX clade are a number of transporters that are genetically linked to the 2,7- anhydro-Neu5Ac oxidoreductase NanOx/NanY (Fig. 1) and in some cases an IT-sialidase, one such transporter being the SP1328 protein from *S. pneumoniae* TIGR4 [53] (Fig. S7). While this suggests that they may function in the uptake of 2,7-anhydro-sialic acid rather than Neu5Ac [52, 53], they occur within a wider range of closely related Neu5Ac transporters including the characterised *C. difficile* protein (Fig. 4). We noted that the IT-sialidases associated with these NanOx-linked SiaX proteins are orthologues of *S. pneumoniae* NanC (Fig. S7), which has as its primary product Neu5Ac2en (N-acetyl-2,3-dehydro-2-deoxyneuraminic acid) [60–62], this being another oxidised form of Neu5Ac found in nature. As the IT-sialidase acts upstream of the transporter, and the NanOx/NanY proteins can also efficiently convert Neu5Ac2en to Neu5Ac [52], we hypothesise that NanOx-linked SiaX transporters might be able to take up this substrate too. Unlike the characterised anhydro-Neu5Ac transporters of the ST7 and ST1 types, which belong to easily distinguishable phyletic groupings (Fig. 3, Fig. S1), and take up only oxidised sialic acid and not Neu5Ac [18, 52, 53], SiaX transporters are all closely related with each other within a single clade regardless of their predicted specificty, and this raises questions as to what might determine the substrate specificity of individual SiaX proteins [22, 47]. Strikingly, all the residues forming the known sialic acid-binding site in the structure of *P. mirabilis* SiaT (Fig. S8) are conserved across the SiaT/X/F proteins, which does corroborate their single origin, but does not help gain insights into their exact substrate specificities. In the absence of detailed structural-functional information about SiaX transporters, their specificity towards Neu5Ac/anhydro-Neu5Ac remains unknown, but it is possible that some of them can transport multiple forms of sialic acid, as reported for other types of sialic transporters. For instance, the ST2-SBP SiaP can bind both Neu5Ac and Neu5Ac2en (though the latter with considerably lower affinity) [14, 35], and the ST1 anhydro-sialic acid transporter NanX has been reported to take up Neu5Ac2en besides 2,7-anhydro-Neu5Ac [52]. These cases provide precedents of sorts for versatile binding sites that recognise different forms of sialic acid.

### Identification of ST5 sialic acid transporters in pathogenic *Spirochaetes*

Using our transporter-led approach has also uncovered more fundamental insight into the sialic acid biology of the pathogenic *Spirochaetes*, organisms where sialic acid transport genes have been previously identified only once (namely, ST2/*siaPQM* genes in *Treponema brennaborense* [33]; Fig. S4). The SiaT clade of ST5 transporters also includes a small sub-clade of highly similar transporters that occur in some species of the Spirochaete genus *Brachyspira* (Fig. 4). *B. hyodysenteriae*, *B. pilosicoli* B2904, and *B. pilosicoli* WesB are pathogens of pigs, birds, and humans, respectively, and have recently been reported to catabolise Neu5Ac/Neu5Gc and adhere to sialic-acid rich mucin glycoproteins [63, 64]. By looking for homologues of SiaT transporters in these bacteria we identified a full complement of sialocatabolic genes in two *B. pilosicoli* strains (Fig. S7), and complete sets of *nan* genes scattered over different loci in the other species including *B. hyodysenteriae* (Fig. S7 and S11), which is consistent with these species’ reported use of sialic acid as carbon source (69). *Brachyspira* spp., *B. pilosicoli* WesB and *B. hampsonii* NSH-16 [65] had satellite loci carrying extra sialic acid transporter genes, specifically a second similar *siaT* gene in WesB (Fig. 4; Fig. S7) and genes for the ST7 2,7-anhydro-sialic acid transporter, SatXYZ (formerly SAT2, Table 1), linked to *nanOx/nanY*, in *hampsonii* (Fig. S11).

### The SiaF clade of ST5 includes SSS transporters fused to sialic acid mutarotases

The third clade of ST5 transporters identified in this analysis, SiaF, is present in a diverse group of bacteria belonging to the phyla Bacteroidetes, Planctomycetes, and Verrucomicrobia (Fig 4). This phylogenetically heterogenous clade is distinguished by a common, hitherto unprecedented feature in that they all contain N-terminal fusions to the NanM Neu5Ac mutarotase protein domain [66] (Fig. S7). The NanM domain carries a predicted leader peptide for its translocation across the membrane, consistent with the periplasmic localisation of unfused NanM in *E. coli* [66] and the N_out_-C_in_ topology of the 13 TMH SSS domain [47] (Table 1), but its 6-bladed β-propeller core appears to lack the helical dimerisation hairpin present in the homodimeric *E. coli* enzyme (see Methods), suggesting that within the fusion the NanM domain is monomeric, just like the attached SSS moiety [22, 47]. The architecture of these fusion proteins is consistent with the proposed role of NanM to act upstream of sialic acid uptake to provide a faster supply of anomerically correct substrate (β-Neu5Ac) to the transporter [66]. Many SiaF transporters are encoded in *nan* clusters (Fig. 2 and S7) with some being the only identified sialic acid transporters, while others co-occur with orthologues of the ST4 MFS transporter (e.g., in *Phocaeicola* – ex *Bacteroides*, *vulgatus*; Fig. S7). Very few exceptions are found outside the above phyla, and all are limited to γ-Proteobacteria species such as *Pseudoalteromonas haloplanktis* (Fig. S7), where these transporters had been identified previously, but not as fusion proteins [13].

The experimental evidence for the function of these SiaF transporters is very scarce. To our knowledge, there are only two cases where sialic acid metabolism has been investigated (to a degree) in bacteria that bear *siaF* genes, namely the Bacteroidetes *Alistipes timonensis* JC36 and *Capnocytophaga canimorsus* [67, 68]. While the best-understood aspect of sialometabolism in *C. canimorsus* is the involvement of the sialidase SiaC in growth [68], in the case of *A. timonensis* we have evidence of sialic acid acquisition in the form of the stimulating effect that exogenous Neu5Ac has on growth in liquid culture [67]. As SiaF is the only predicted sialic acid transporter in this latter bacterium (Fig. S7), this provides good preliminary evidence for SiaF proteins to function in sialic acid uptake, but more detailed investigation is required to clarify the function of these proteins and to elucidate the role of the NanM-ST5 fusion.

### The ST4 (NanZ) MFS transporters are distinct from ST1 (NanT/X) proteins and occur in Bacteroidetes and other diverse gut commensals

A well-studied gut bacterium that is known to rapidly consume host-derived sialic acid is *Bacteroides fragilis* (68,29,30). As mentioned in the introduction, the transporter from this bacterium is a known MFS transporter and was perhaps understandably called NanT, as the canonical *E. coli* NanT (ST1) is also an MFS transporter. Our analysis, however, places the *B. fragilis* MFS transporter in an evolutionary distinct family, which we have named ST4 with the transporter renamed NanZ (Fig. 3A; Table 1). ST4 proteins are also found in the phyla Planctomycetes and Verrucomicrobia, forming a distinct clade (Fig. S1-3), and are not seen in any other bacteria. In *B. fragilis* we identified three *nanZ* genes (*BF1633*, *BF3607*, and *BF3947*), encoding proteins of ca. 80% identity yet only the *BF1633* gene product was previously described as a sialic acid transporter (Table 1), seemingly accounting for all sialic acid acquisition in this bacterium under the conditions tested [30]. Unlike the close orthologue from the oral pathogen *Tannerella forsythia* [31], neither *B. fragilis* NanZ (BF1633) nor its two BF3607 and BF3947 paralogues have been studied in an heterologous host, so it is not known whether these transporters are functionally different or whether the redundancy reflects use in different environmental conditions. It is notable that other bacteria such as the sialic acid-utilising commensal *Muribaculum intestinale* [67] also have multiple *nanZ* genes (Fig. S3).

Remarkably, we also found two instances of NanZ sequences in the important gut bacterium *Akkermansia muciniphila* and the closely related species, *A. glycaniphila*, where the ST4 transporter is again fused to a NanM-like domain (Fig. S3), as we have just described for the SiaF proteins of ST5. Similarly, the N-terminal mutarotase domain carries a leader peptide for translocation to the extracytoplasmic space, while the transporter moiety, which in NanZ proteins is normally made of a predicted 12 TMH N_in_-C_in_ core (Table 1), here possesses an extra N-terminal TMH to adjust the topology accordingly. In both *Akkermansia* species, the corresponding genes are part of a separate locus outside the *nan* cluster (Fig. S3). Whereas some studies reported that *A. muciniphila* can release but not consume sialic acid [70], others reported growth stimulation by Neu5Ac for the same strain in a complex medium [67]. It is therefore possible that this fusion protein may act as a Neu5Ac transporter in *Akkermansia* species, under specific growth conditions, although experimental confirmation of its function is warranted. To our knowledge, this is the first identification of a candidate sialic acid transporter for these species, and this also completes the mapping of sialocatabolic genes among Planctomycetes and Verrucomicrobia [5, 9, 13].

With regard to the ST1 family, which contains NanT and NanX proteins, we observed overlapping yet distinctive distributions for the two clades, with NanT orthologues primarily found in Enterobacteria and Actinobacteria, and NanX orthologues occurring in different orders of γ-Proteobacteria (Fig. S1-2). To our knowledge this is the first report of ST1 sialic acid transporters occurring outside the γ-Proteobacteria. Notably, all NanX transporters are genetically linked to the NanOx/NanY oxidoreductase in these novel genotypes (Fig. S2), indicating that they might function as 2,7-anhydro-sialic acid transporters in these organisms too. NanT and NanX can also be distinguished as they contains a different numbers of TMHs, 14 in NanT [26] and 12 in NanX (Table 1), which as previously proposed [4] might account for the different substrate specificity of the transporters.

### Expanded distribution of ST2 (SiaPQM) TRAP transporter in bacterial pathogens

So far, we have discussed sialic acid transporters from the ST1, ST4, and ST5 families, accounting for MFS and SSS transporters; from these, the 4 remaining ST types that feature characterised transporters (ST2, ST3, ST6 and ST7) are distinct in that they use an SBP in their mechanism, which usually correlates with higher affinity transport than that conferred by a classical symporter.

For the ST2 proteins, which are TRAP transporters, we expanded their distribution from exclusively Gram-negative organisms (primarily γ-Proteobacteria and Fusobacteria, but also the Spirochaete *T. brennaborense* [33]) to include Firmicutes, which are here reported for the first time (Fig. S4). Recent evidence supports an important function of ST2 proteins in bacterial vaginosis, as the pathogen *Fusobacterium nucleatum* is able to use sialic acid liberated by other members of the vaginal microbiota to improve its colonisation of this niche [19]. The transporter used by *F. nucleatum* was originally identified as a SiaPQM system in early studies on ST2 sialic acid transporters [14], and confirmation of its role in sialic acid acquisition and growth was obtained by deleting the gene for the fused membrane component, *siaQM* (called *siaT* in [19]; Table 1). Interestingly, we found ST2 transporters in *nan* clusters in two other species of *Fusobacterium*, namely, *F. ulcerans* and *F. mortiferum* (Fig. S4), and recent data suggest that *F. mortiferum* can also consume free sialic acid similar to its cousin *F. nucleatum* [19, 71].

### The ST3 (SatABCD) ABC transporters are widespread in the Actinobacteria

Our analysis of the ST3 proteins (Fig. S5-S6) revealed a significant change in our understanding of the origins of this ABC transporter. While the original SatABCD system was discovered and characterised in the Gram-negative bacterium *Haemophilus ducreyi*, with other examples pointed out in related *Pasteurellaceae* [38], it is now clear that their origin lies in the Actinobacteria (Fig. S5-S6). Experimental support for this assertion comes from characterisation of ST3 transporters in both *Corynebacterium glutamicum* and *Bifidobacterium breve*, where the genes encoding the ST3 system have been disrupted with resulting loss of growth on Neu5Ac [39, 72]. The same genes are in the characterised *nan* cluster of *Bifidobacterium longum* subsp. *infantis* [40]. Also, the vaginal pathogen *Gardnerella vaginalis* ATCC 14019, which is a known sialidase-positive Neu5Ac consumer, contains an ST3 family transporter very similar to the characterised Bifidobacterial system [9], which is thus likely the route of sialic acid uptake by this important pathogen [73]. A ST3 system is also seen in the Actinobacterium *Micromonospora viridifaciens*, a non-pathogenic bacterium identified as a producer of high levels of an inducible sialidase activity [74]. We describe the full *nan* cluster from this bacterium for the first time (Fig. S6), which contains the ST3 transporter and catabolic genes as well as a gene for a likely sialic acid-responsive transcription factor. Also, this cluster contains the structural gene for the well-studied GH33 family sialidase [75]. Hence, it is highly likely that sialic acid utilisation plays a role in the biology of this bacterium in the soil as has been suggested for the related Actinobacterium *C. glutamicum* [39, 76]. Our analysis now suggests that the small *Pasteurellaceae* clade of ST3 sialic acid transporters originated by horizontal gene transfer (HGT) from Actinobacterial clusters.

### ST6 (SatEFG) transporters in Spirochaetes and chemotaxis

The distribution of ST6 ABC transporters identified orthologues primarily in Firmicutes (Fig. S9), including the known pathogen *Streptococcus pneumoniae* and other important species of *Streptococcus* [16, 77]. A ST6 transporter is also present in the sialic acid-utilising commensal *Ruthenibacterium lactatiformans* [67] (Fig. S10). We renamed this transporter SatEFG to avoid confusion with ST3 transporters (Table 1); the ATPase component, which is normally provided by the multitasking *msmK* gene [44], should be called SatH when encoded by a dedicated gene in the same cluster (Fig. S10). These transporters are also seen in a small number of Spirochaetes including *T. brennaborense* (Fig. S10); however, we also discovered a few instances of a novel genetic linkage between the sole SBP component (SatE) of a ST6 system and a methyl-accepting chemotaxis protein (MCP [78]) (Fig. S10). These instances all occur in some *Treponema* species including the ‘Red Complex’ [79] pathogen, *T. denticola* (Fig. S10). As reported by others [79, 80], we could not find orthologues of any *nan* catabolic and/or other sialic acid transporter genes in *T. denticola* (unlike in *T. brennaborense*, which bears ST2, ST6, and *nan* genes; Fig S4-S10), though there is some evidence for Neu5Ac consumption by *T. denticola* [80]. We propose that the orphan ST6-SBP/SatE orthologue in *T. denticola* and other *Treponema* species may play a role analogous to that of other SBPs as part of a sensory apparatus for the chemotactic response to small molecules (in this case, sialic acid), which is mediated by physical interaction with MCPs [81].

### The ST7 (SatXYZ) and ST8 (NanG) families contain additional transporters for 2,7-anhydro-Neu5Ac

Of the final two STs, ST7 includes the recently discovered class of ABC transporters with specificity for 2,7-anhydro-Neu5Ac, exemplified by the characterised system from *Ruminococcus gnavus* [18, 51]. We propose that these are called SatXYZ (Table 1), while the ATPase component should be called SatW when encoded by a dedicated gene in the same cluster (as in e.g., *Haemophilus haemoglobinophilus*, Fig. S11). Novel examples of ST7 transporters are those found in the Spirochaete *B. hampsonii* NHS-16, a porcine pathogen [65], and the human oral pathogen *Streptococcus sanguinis* SK36, but intriguingly ST7 genes also occur in the environmental Actinobacterium *Beutenbergia cavernae* (Fig. S11), suggesting a yet unidentified role in the environment (as argued above for ST3 systems).

The final family, ST8, encompasses MFS transporters of the GPH type from within the MFS_2 Pfam family, which is distinct from the MFS_1 family featuring ST1 (NanT/X) and ST4 (NanZ) transporters (Fig. 3). We propose to call ST8 transporters NanG (for GPH). The only ST8 transporter identified prior to this work was found in some strains of *L. salivarius* [52, 53], and here we discovered more orthologues through the conserved linkage to the NanOx/NanY oxidoreductase, and in some cases also to an IT-sialidase of the type of *S. pneumoniae* NanB, which has 2,7-anhydro-Neu5Ac as its the primary product [61] (Fig. S12). These genetic links suggest that ST8 transporters might specialise in the uptake of 2,7-anhydro-Neu5Ac, but this will need to be confirmed experimentally. The distribution of the newly identified ST8/NanG transporters [53] strongly suggests that this family might have evolved recently, as orthologues primarily occur in a very small number of Firmicutes and are highly similar (>70% aa identity) proteins (Fig. S12).

## DISCUSSION

This *in silico* analysis provided novel insights into the evolution of bacterial sialic acid transporters, complementing the comprehensive work on the downstream catabolic enzymes [9, 13, 33, 34]. Using phylogeny as the basis for classification, we found that sialic acid transport has evolved no fewer than eight times from four major superfamilies of transporters (Table 1; Fig. 5). The most diversity was found within the ST5/SSS transporters despite them likely having a single origin, and this family provided the first examples of fusions of transporters to other sialic acid metabolism related proteins (SiaF). The only time the phylogenetic distribution of sialic acid transporter genes has been considered previously was in the comprehensive analysis of the distribution of the NanA enzyme, where transporter families were mapped onto the NanA phylogeny [9], using the broader classification to four families, namely SSS, MFS (NanT), ABC, or TRAP [9]. The detailed and comprehensive study of human gut microbiome organisms by Ravcheev & Thiele [82] used the same four families for general classification, giving all transporters identified, however, arbitrary new names (Table 1). This unique transporter-focussed study now defines 8 sialic acid transporter families (ST1-8) and proposes a unified naming scheme for these systems to reduce the significant confusion current in the community, and to provide a clear naming framework as new bacteria are identified that use sialic acid uptake in their biology. It is also the first study to include a classification of the recently discovered sialic acid transporters with specificity for the anhydro-form released by IT-sialidases and requiring the NanOx/NanY enzyme for utilisation (Fig. 5). Our study now reveals that this latter specificity is present in at least 4 of the 8 STs, suggesting that this is an ancient adaptation that is used by diverse bacteria to consume anhydro-sialic acid released in their environment (Fig. 5).

**Figure 5.**
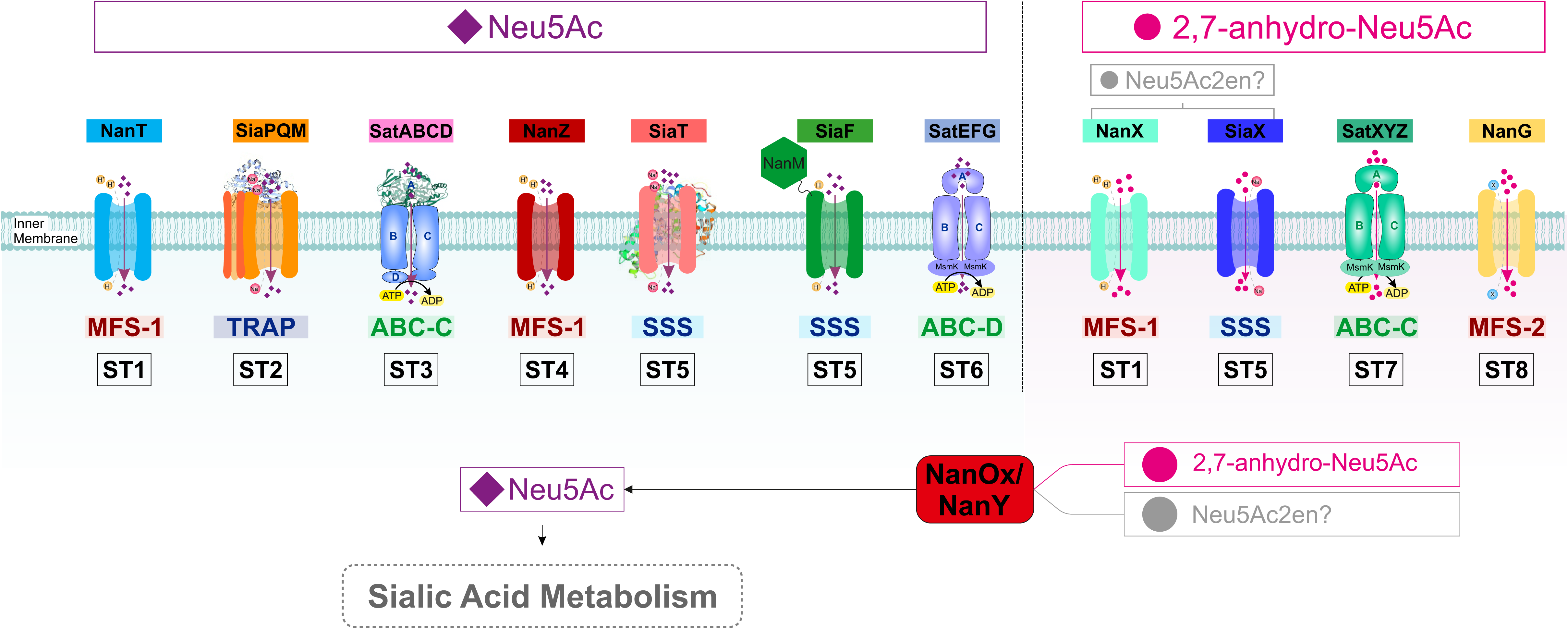
Sialic Acid Transporters ST1-ST8. Newly classified sialic acid transporter families (Table 1) are shown. ST groups are separated by substrate specificity for Neu5Ac or 2,7-anhydro-Neu5Ac/Neu5Ac2en. Substrate specificities are discussed in the Text and summarised in Table 1.

Our analysis identified for the first time candidate sialic acid transporters in Planctomycetes and Verrucomicrobia [5, 13], and expand the repertoire of sialic acid transporters in Spirochaetes, from one type of transporter [33] to four. Recent research has provided evidence for Neu5Ac utilisation by organisms expressing these novel transporters [63, 64, 67], thus lending support to our assignment. The recent observation by Pereira and colleagues that *A. muciniphila*’s growth was stimulated by the addition of Neu5Ac *in vitro* [67], supports a potential biological role for the sialic acid transporters identified in this work. Heterologous expression of these novel candidates will help determine their role in sialic acid transport, as done for the original SiaT [29].

One behaviour being better understood in the function of bacteria in the human gut microbiome is the cross-feeding between commensals and its exploitation by pathogens [15, 82]. One established example concerns the well-characterised common gut commensal *Bacteroides thetaiotaomicron* VPI-5482, which is able to liberate sialic acid by using its sialidase, but not to consume the released Neu5Ac [69], and is therefore thought to act “altruistically”. In keeping with these results, it has been stated that this bacterium lacks the sialic acid catabolic genes [15]. With regard to this latter statement, we note however from the analysis presented in this paper that *B. thetaiotaomicron* does contain an ST5 family transporter (Fig. 4), which in fact retains all the residues of the Neu5Ac-binding site identified in *P. mirabilis* SiaT (Fig. S8). This gene, *BT_2813*, sits next to *BT_2814*, which encodes a NanA-like protein, and in some other related Bacteroidetes such as *Cyclobacterium marinum* (Fig. 4) the same two genes are linked to a sialidase gene. Also at a separate locus there is a *nanE-II* gene, *BT_3605*, linked to orthologues of the *nanOU* outer membrane sialic acid transporter genes from *B. fragilis* [83], suggesting that *B. thetaiotaomicron* actually has the required complement of gene functions to take up and catabolise some form of sialic acid or even Neu5Ac. While the evidence for sialic acid cross-feeding in the gut is clear, our findings at least suggest that *B. thetaiotaomicron*’s capacity for sialic acid utilisation should be checked in more detail (e.g., by heterologous expression of the transporter gene [48]) before the idea that this bacterium cannot use sialic acid at all is set in concrete [15].

Our discovery of protein fusions between the sialic acid mutarotase, NanM, and two unrelated types of sialic acid transporters (ST4 and ST5) is the first ever report of a covalent coupling between a metabolic enzyme and a transporter in sialic acid biology. It is intriguing that, in both of these independent instances of fusion, the fusion takes place at the N-terminus of the transporter, rather than at the C-terminus where is most frequent in bacteria [84, 85]; however, the extracytoplasmic positioning of the NanM moiety makes mechanistic sense and can be understood thinking of the location of NanM as a periplasmic enzyme. The biological function of NanM in the periplasm has been proposed to be to help bacteria acquire more rapidly the transported anomer of Neu5Ac, which is the beta-anomer [66] (Fig. 5). In this line of reasoning, it makes teleological sense for a transporter to recruit NanM as an integral domain located in the periplasm, as this domain would feed β-Neu5Ac directly into the transporter as opposed to releasing it in solution where diffusion would limit its acquisition by the transporter.

Remarkably, our work uncovered a potential new biological role for a sialic acid transporter component, ST6-SBP SatE, in *T. denticola*. Together with *Porphyromonas gingivalis* and *T. forsythia*, *T. denticola* forms the so-called Red Complex, a consortium of microorganisms that underlie the inflammatory condition known as periodontitis leading to tooth loss [79]. While the role of sialic acid metabolism in periodontal pathogens is still under investigation, both *P. gingivalis* and *T. forsythia* secrete sialidases for the release of free Neu5Ac, which *T. forsythia* can also consume [31, 79]. In the light of our observation, it is tempting to speculate that the suggested SatE-dependent chemotactic response to sialic acid by *T. denticola* might contribute to the establishment of and/or the dynamics within the Red Complex.

We have previously considered the consequences of using different types of transporter mechanisms for sialic acid acquisition and attempted to rationalise this to the biological niche that the bacterium inhibits [12]. Given this larger dataset it is worth considering this again in, for example, the anaerobic environment of the human colon. Bacteria here are growing by fermentation by scavenging dietary and host-derived carbon and energy sources. While one might expect transport to use the most energetically efficient systems in this niche, the reality is that the bacteria that live there use all varieties of transporters mechanisms for sialic acid uptake. This analysis is complicated now by the occurrence of often more than one sialic acid transporter in a single bacterium, which potentially allows the organism to use different transporters under different environmental conditions. The reason to have a Neu5Ac transporter and a separate 2,7-anhydro-Neu5Ac transporter for a gut bacterium appears clear, as both substrates will be liberated from host glycans depending on which sialidases are made by the community, and it does seem advantageous for a species to be able to take up whichever form is available. However, it is not clear why bacteria have multiple Neu5Ac transporters, although again this might relate to individual preferences for other forms of sialic acid, such as Neu5Gc available through the diet. Also, having multiple transporters for Neu5Ac with varying affinity could be a good reason to have apparently redundant systems, as again it would allow the bacterium to scavenge sialic acid most efficiently, if its concentration were changing in the environment.

As microbiome-sequencing projects resolve an ever-growing number of microbial communities in health and disease, the diversity among sialic acid transporters is likely to increase and research is warranted to decipher the functional role and substrate specificity of these important transporters. Our naming system, which matches the 8 phylogenetically different families of transporters, now provides a clear framework to accommodate newly identified members within these families and can be expanded as new sialic acid transporter families are discovered in the future.

## METHODS

### Sialic acid transporter sequences

The criteria used to collate sialic acid transporter sequences were based on experimental evidence and *in silico* analysis of the sialic acid *nan* operon. Using functionally characterised sialic acid transporters as initial queries we searched for homologous proteins based on sequence similarity using BLAST [12, 18], and all hits were validated by reciprocal BLAST searches. For SBP-dependent transporters (Fig. 1), we used the SBP component as search query, as done in [33]. As we added new clades to the phylogenetic trees, we took a heuristic approach and included the new members in both direct and reciprocal searches, and re-validated the hits found in previous rounds. At the onset of these searches, homologous yet non-orthologous hits were included in order to resolve the phylogeny. Assignment of orthologous transporters was aided with the mapping of Neu5Ac-binding residues when these were known from crystallographic and/or mutational studies (Table 1). All organisms where we found candidate orthologues of sialic acid transporters were confirmed for the presence of complete sialocatabolic pathways (encoded within clusters or separate loci), recapitulating the methodology and the results described in [13, 34]. As queries, we used sequences of different organismal origin for the following enzymes: NanA (Neu5Ac lyase), NanK (ManNAc kinase), NanE (ManNAc-6P epimerase), and NanE-II (ManNAc epimerase) (Fig. S1-12). Clusters were then annotated outwards to identify further distinctive sialometabolic genes in the clusters, such as e.g., sialidase genes, NanM (Neu5Ac mutarotase), NagA (GlcNAc-6P deacetylase), and NagB (GlcN deaminase). All sialic acid transporter-bearing organisms were searched for further orthologues of the same or different transporter families. Number of TMH and presence of signal peptides were predicted using TMHMM [86] and SignalP5 [87], respectively. 3D structure prediction for NanM domains was performed with i-Tasser [88] and Swiss-model [89]. Wherever possible, we opted for fully over partially sequenced genomes in order to minimise incomplete genetic information. All sequences were retrieved from NCBI database.

### Phylogenetic analysis

Each archetypal sialic acid transporter was used a search query in the PFAM database. To reduce the sequence dataset within a PFAM family we downloaded the representative proteome RP15 for each PFAM identifier. Unique accession numbers were retrieved from this list and used to download full length proteins from the Uniprot database. Bacterial sequences were filtered from each PFAM family and compiled with our heuristically predicted sialic acid transporters to generate multiple sequence alignments. For large alignments (>500 sequences) sequences were aligned with MUSCLE using default settings. Alignments were manually inspected using Jalview [90]. For sequences <200, alignments were generated using MAFFT with L-INS-I for an accurate amino acid alignment. Phylogenetic reconstruction was performed using IQ-Tree [91]. Phylogeny was inferred by maximum likelihood with automatic model selection to find the best-fit model. Ultrafast bootstrap approximation was used give branch support. Phylogeny was visualised with iTOL [92]. Alignment of selected proteins with crystal structures was made with ESPript [93]. All figures were prepared using CorelDRAW 2020 and PowerPoint.

---

**Figure S1.**
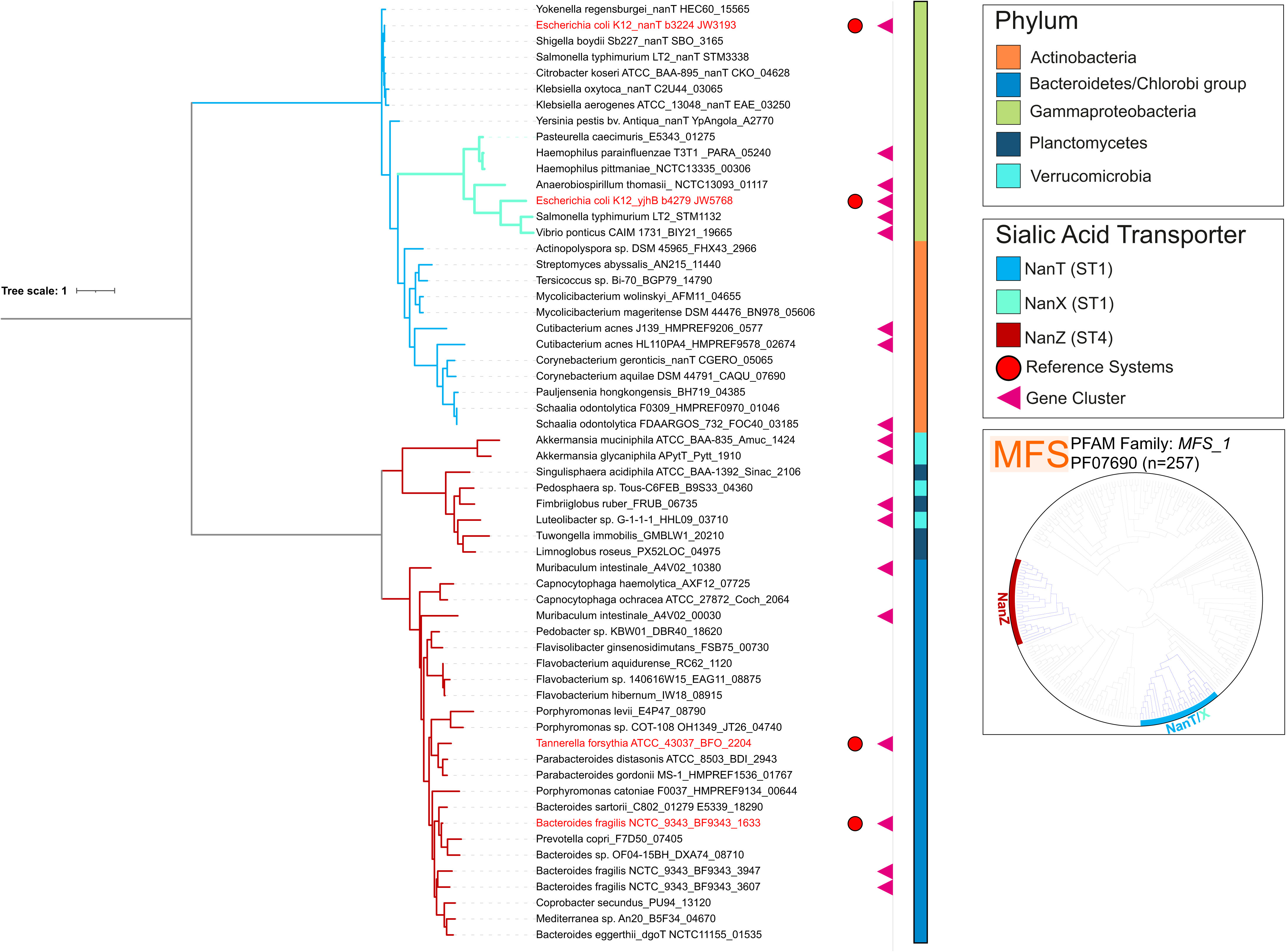
Phylogeny of MFS sialic acid transporters of the ST1 and ST4 families. Maximum likelihood phylogeny of MFS_1 sialic acid transporters that reside within *nan* clusters. ST1 sialic acid transporters clearly separate into two distinct clades, NanT and NanX. This broadly reflects two different substrate affinities where NanX has been shown to transport 2,7-anhydro-Neu5Ac/Neu5Ac2en and NanT selectively transports Neu5Ac (see Text). NanZ is found widely among Bacteroidetes, Planctomycetes, and Verrucomicrobia, and has been shown to transport Neu5Ac in Bacteroidetes (see Text). Reference systems are highlighted in red.

**Figure S2.**
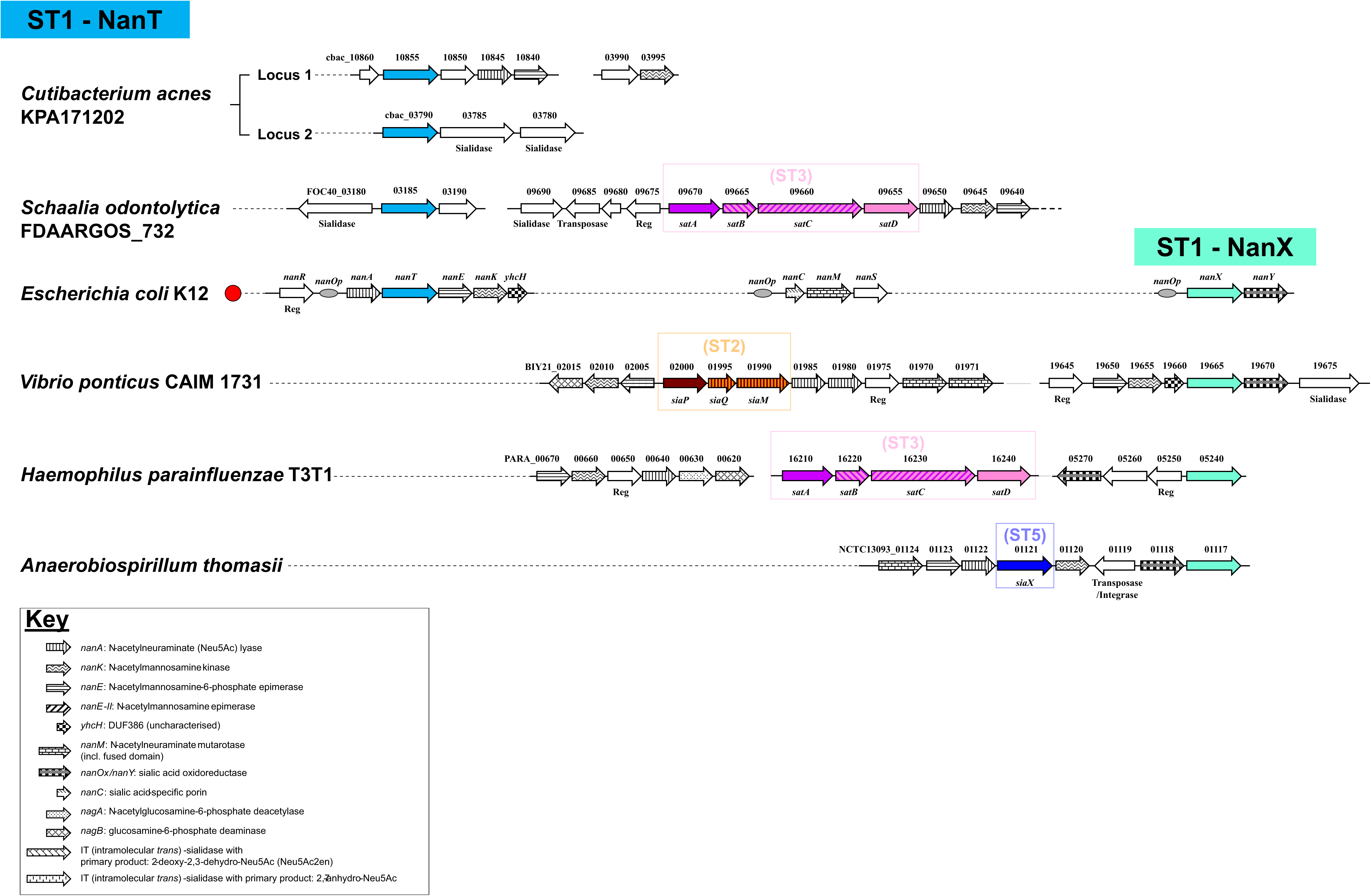
Organisation of ST1 (*nanT/X*) gene clusters in selected bacteria. ST1 transporter genes *nanT* and *nanX* are colour-coded. Catabolic *nan* genes are pattern-coded and described in the key. For clarity other conserved genes recurring in these clusters are not pattern-coded except when emphasising specific genetic links. The function of other notable genes is reported (without pattern) if relevant for cluster annotation or to exemplify instances of potential HGT. Reference gene clusters are highlighted with a red circle. *nanOp* operators highlighted in the *E. coli* ST1 loci emphasise the occurrence of a single NanR regulon in this organism. Reg: regulator gene; SBP: solute-binding protein; TMD: transmembrane domain; NBD: nucleotide-binding domain (ATP-ase protein); HGT: horizontal gene transfer.

**Figure S3.**
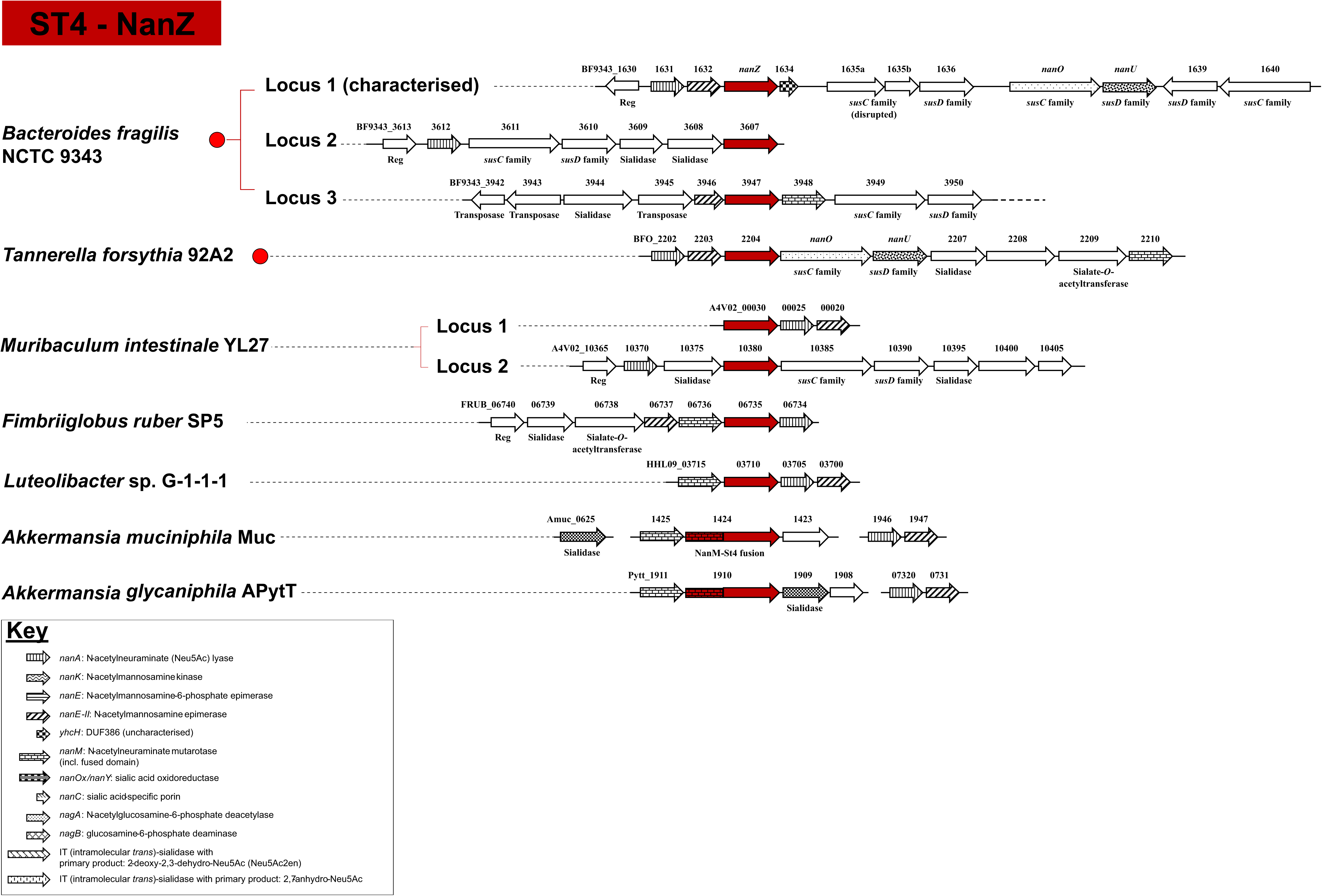
Organisation of ST4 (*nanZ*) gene clusters in selected bacteria. NanZ occurs in gene clusters with other catabolic *nan* genes required for sialic acid metabolism. *nanZ* is colour-coded, catabolic *nan* genes are pattern-coded and described in the key. For clarity other conserved genes recurring in these clusters are not pattern-coded except when emphasising specific genetic links. The function of other notable genes is reported (without pattern) if relevant for cluster annotation or to exemplify instances of potential HGT. Interestingly, there are two novel examples of a NanM-NanZ mutarotase-transporter fusions found in *Akkermansia* spp. (NanM-like domains are indicated by the appropriate pattern). Reg: regulator gene; SBP: solute-binding protein; TMD: transmembrane domain; NBD: nucleotide-binding domain (ATP-ase protein); HGT: horizontal gene transfer. Reference gene clusters are highlighted in red. SusCD: outer membrane protein complex for glycan acquisition made of a TonB-dependent transporter (SusC) and an extracytpolasmic lipoprotein (SusD). NanOU is an experimentally confirmed sialic acid-specific SusCD-family complex [83]. We did not establish orthologous groups for all other, more divergent SusCD-family complexes in this figure.

**Figure S4.**
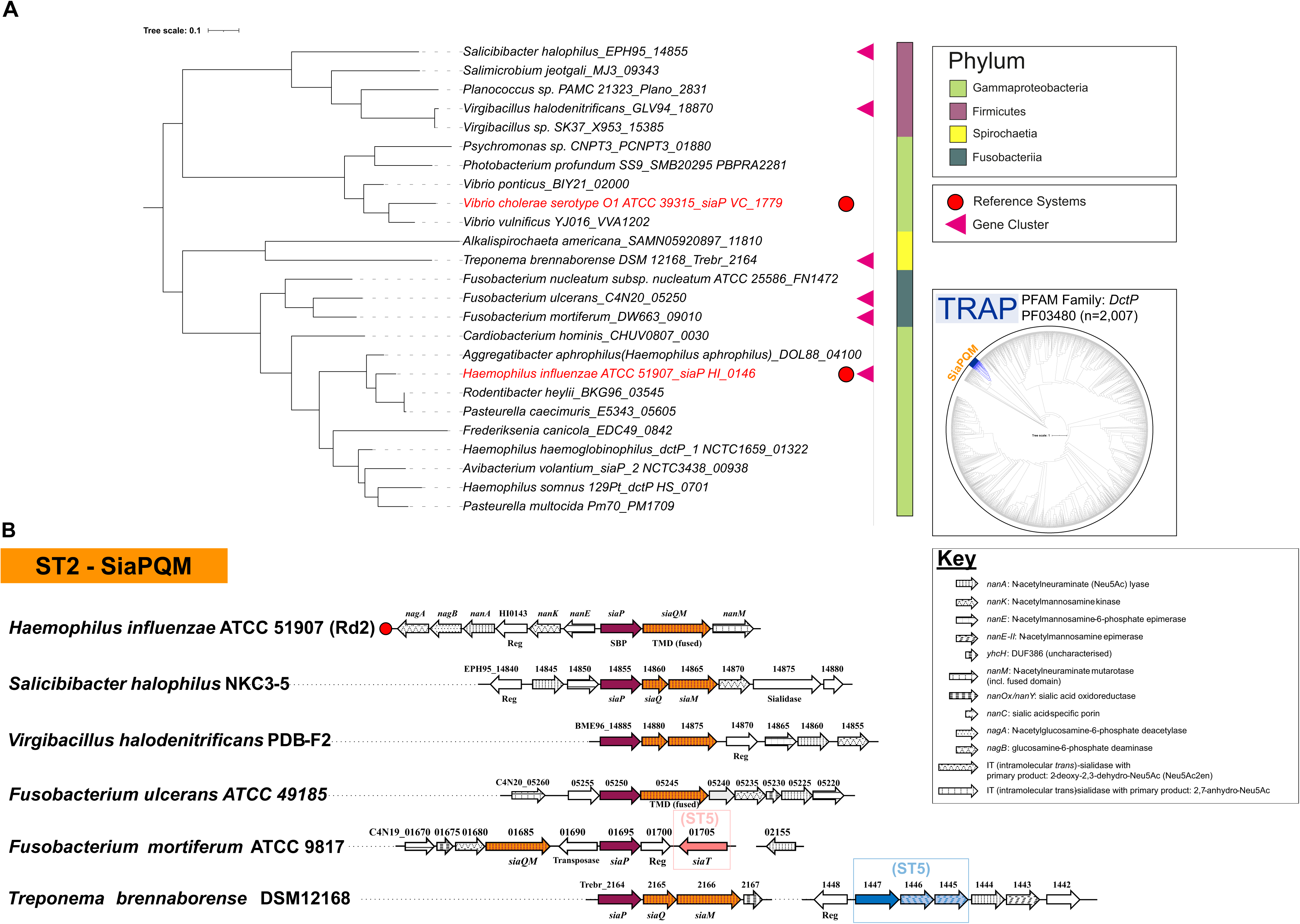
Phylogeny of the ST2-SBP, SiaP, and organisation of ST2 (*siaPQM*) gene clusters in selected bacteria. **(A)** Maximum likelihood phylogeny of sialic acid-binding extracytoplasmic protein, SiaP. Reference organisms are highlighted in red. **(B)** Genetic organisation of *siaPQM* transporter genes within *nan* clusters. The figure emphasises the finding of ST2 genes in Firmicutes, reported here for the first time [33], but also includes *Fusobacterium* spp. and *T. brennaborense* discussed in the Text. ST2 transporter genes are colour-coded, patterns are used to distinguish among different subunits. Catabolic *nan* genes are pattern-coded and described in the key. For clarity other conserved genes recurring in these clusters are not pattern-coded except when emphasising specific genetic links. The function of other notable genes is reported (without pattern) if relevant for cluster annotation or to exemplify instances of potential HGT. Reference gene clusters are highlighted in red. Reg: regulator gene; SBP: solute-binding protein; TMD: transmembrane domain; NBD: nucleotide-binding domain (ATP-ase protein); HGT: horizontal gene transfer.

**Figure S5.**
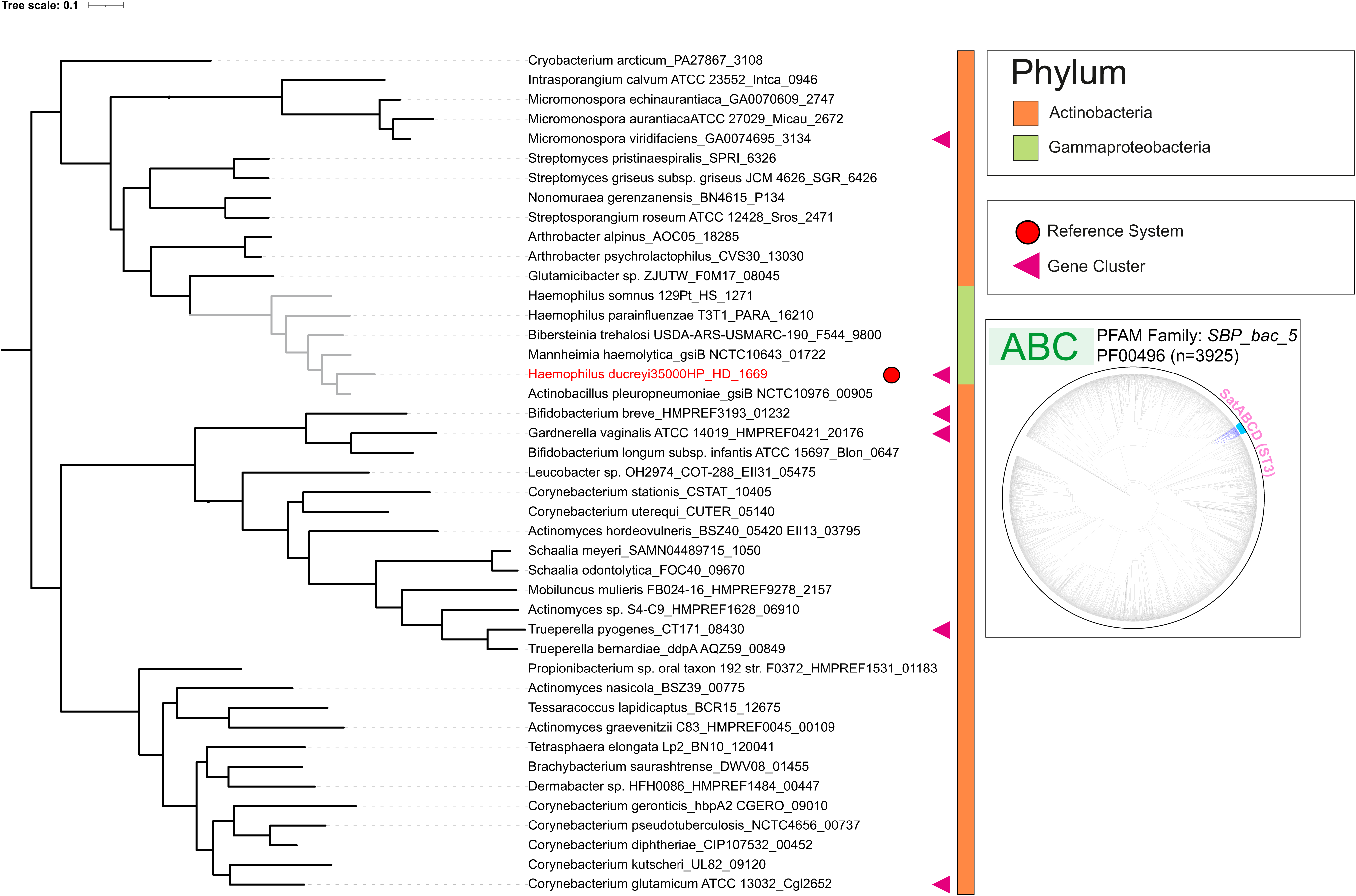
Phylogeny of the ST3-SBP, SatA. Maximum likelihood phylogeny of sialic acid-binding protein component (SatA) of the ST3 ABC transport system, SatABCD. Reference organisms are highlighted in red. Note how sequences from *Pasteurellaceae* form a small clade (in grey) within a diverse family of ST3 transporters that is otherwise exclusive to Actinobacteria.

**Figure S6.**
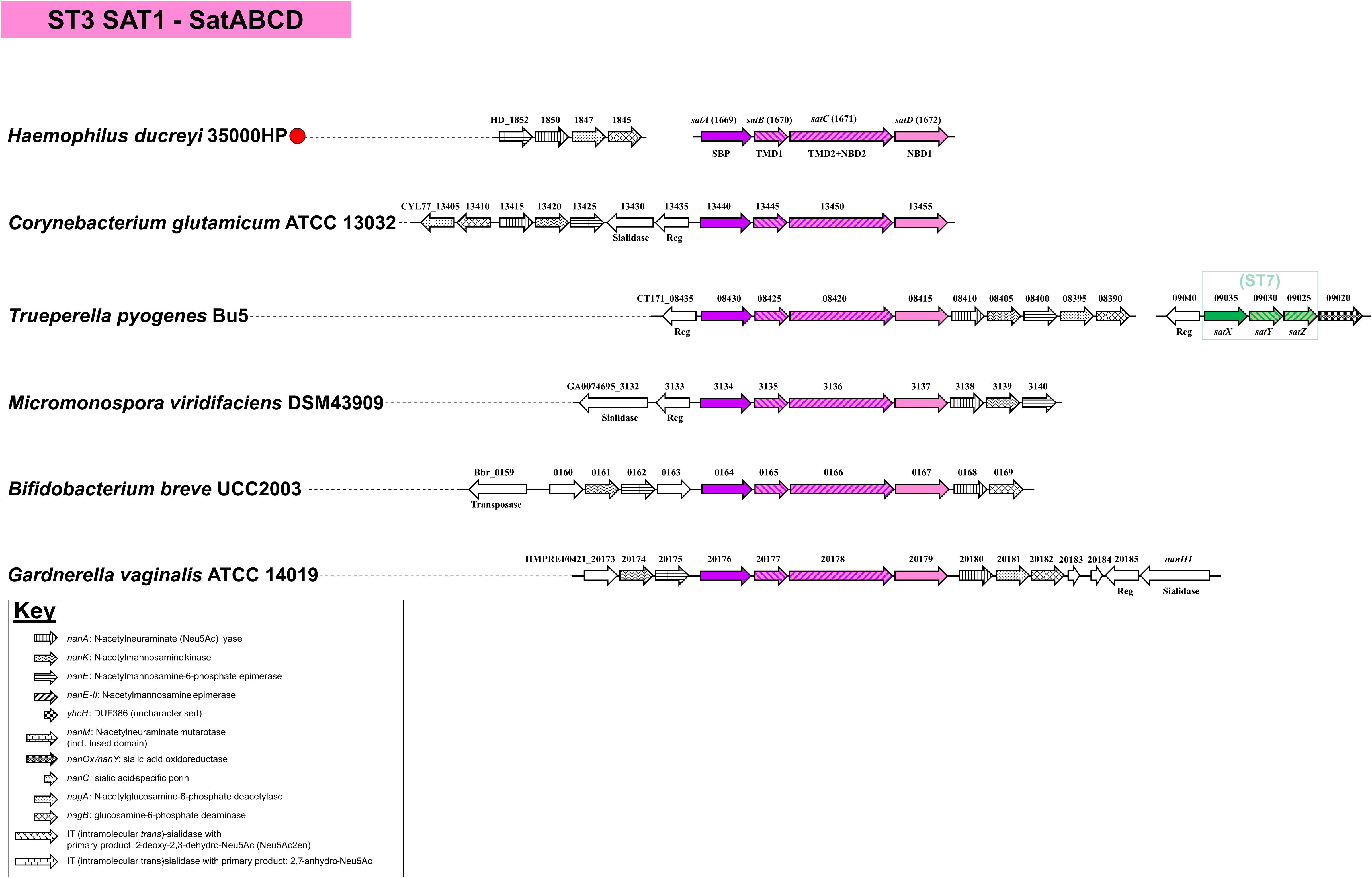
Organisation of ST3 (*satABCD*) gene cluster in selected bacteria. ST3 transporter genes are colour-coded, patterns are used to distinguish among different subunits. Catabolic *nan* genes are pattern-coded and described in the key. For clarity other conserved genes recurring in these clusters are not pattern-coded except when emphasising specific genetic links. The function of other notable genes is reported (without pattern) if relevant for cluster annotation or to exemplify instances of potential HGT. Reference gene clusters are highlighted with a red circle. Note that no *nanK* orthologues occur in *H. ducreyi* ([38] and this work), and genes encoding for ManNAc kinase function are yet to be identified (see also Fig. 2). Reg: regulator gene; SBP: solute-binding protein; TMD: transmembrane domain; NBD: nucleotide-binding domain (ATP-ase protein); HGT: horizontal gene transfer.

**Figure S7.**
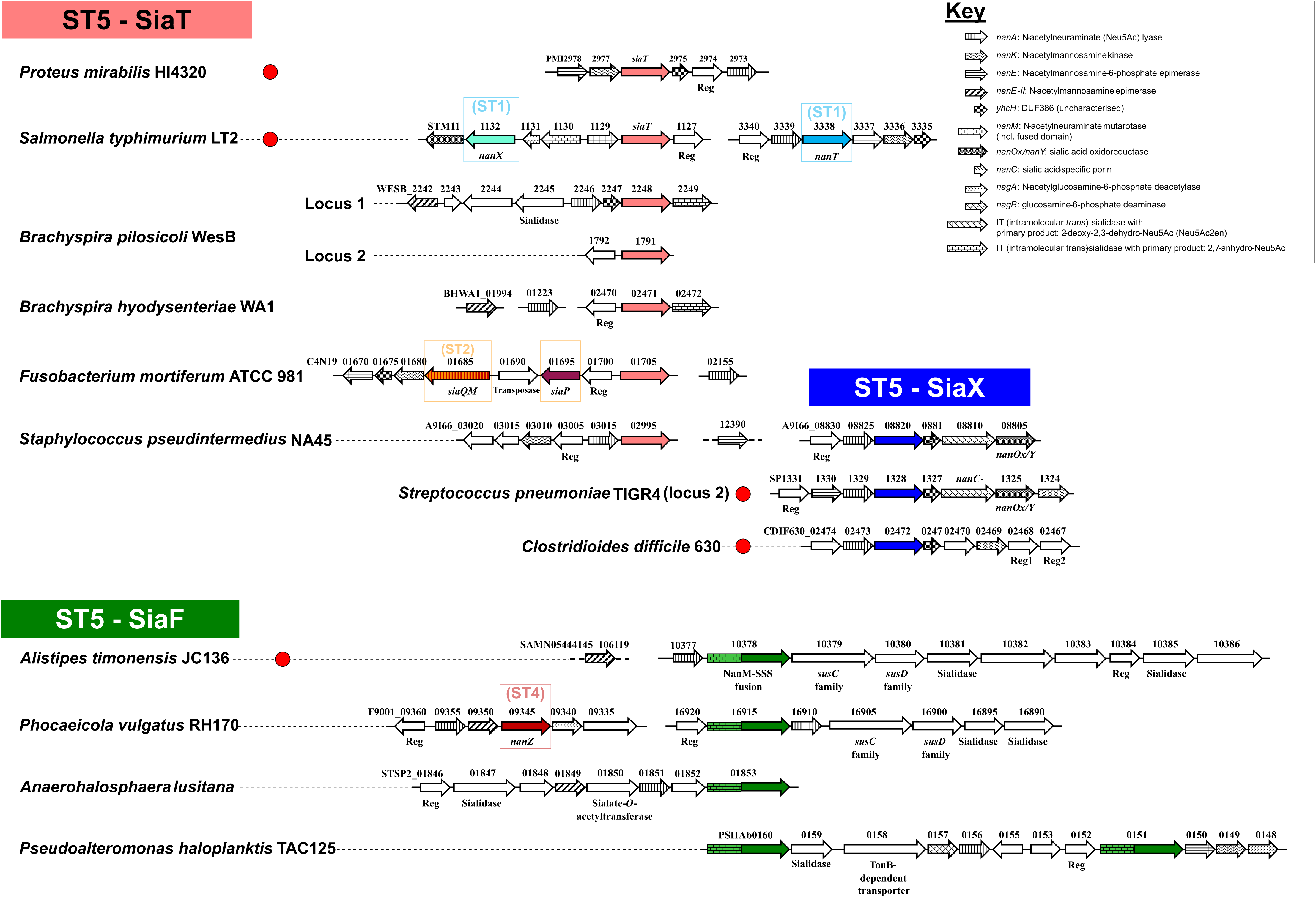
Organisation of ST5 (*siaT*/*siaX*/*siaF*) gene clusters in selected bacteria. Based on phylogeny, the ST5/SSS transporters can be resolved into three distinct clades (Fig. 4). The different transporter genes are colour-coded based on their clade within the ST5 phylogeny. Catabolic *nan* genes are pattern-coded and described in the key. For clarity other conserved genes recurring in these clusters are not pattern-coded except when emphasising specific genetic links. The function of other notable genes is reported (without pattern) if relevant for cluster annotation or to exemplify instances of potential HGT. Reference gene clusters are highlighted by a red circle. SiaT clusters are widespread in bacteria and all characterised transporters from these clusters have been shown to transport Neu5Ac. The *siaT* sialic acid utilisation cluster of *B. pilosicoli* B2904 (not shown) is identical to locus 1 of strain WesB, but locus 2 is present only in WesB. The SiaX clade is common among Firmicutes and includes both Neu5Ac and potential anhydro-Neu5Ac transporters. As for the Neu5Ac transporters of this clade, there is an example of demonstrated Neu5Ac specificity for the *C. difficile* transporter. Candidate anhydro-Neu5Ac transporters are genetically linked to anhydro-Neu5Ac metabolic enzymes, namely the oxidoreductase NanOx/NanY and an IT-sialidase, as seen here in *S. pneumoniae* TIGR4 and *S. pseudintermedius* NA45 (the latter organism also possesses a *siaT* orthologous gene at a different locus). The SiaF clade, occurring near-exclusively in Bacteroidetes and Planctomycetes/Verrucomicrobia, feature proteins that are all fusions between the sialic acid mutarotase NanM (pattern-coded accordingly) and a ST5 transporter. SusCD: outer membrane protein complex for glycan acquisition made of a TonB-dependent transporter (SusC) and an extracytpolasmic lipoprotein (SusD).

**Figure S8.**
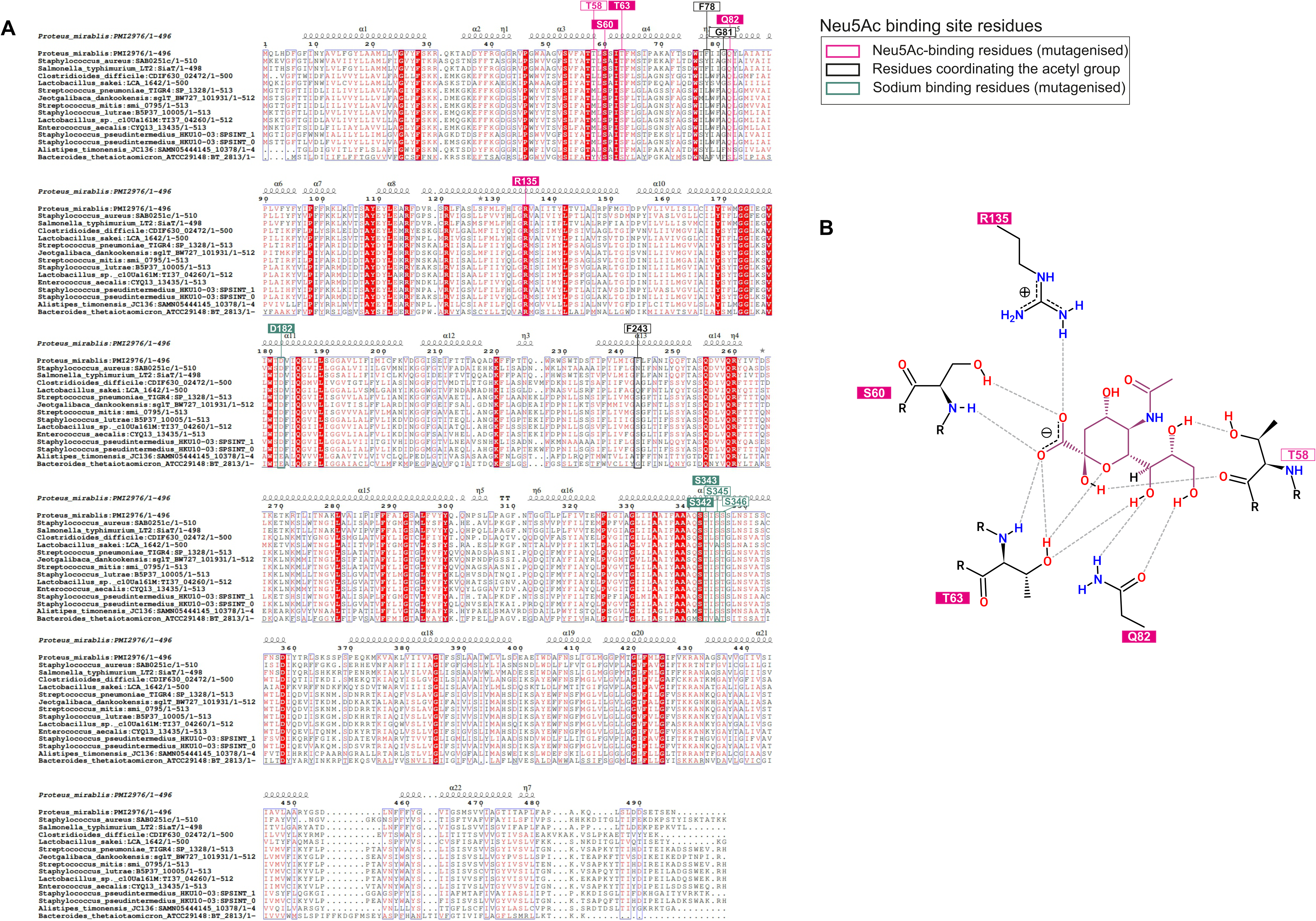
The sialic acid binding-site is conserved across ST5 sialic acid transporters. **(A)** Clustal alignment of ST5 sequences highlighting the Neu5Ac-binding residues (red and black boxes) and selected Na^+^-binding residues (green boxes) in PmSiaT [47]. Residues in red and black boxes have been subjected to mutagenesis, with deleterious effects only seen for residues in block colour [47]. The alignment, visualised with ESPript (see Methods) for mapping to the PmSiaT structure (Table 1), includes all candidate anhydro-Neu5Ac transporters of the SiaX clade listed in Fig. 4, all characterised SiaT and SiaX Neu5Ac transporters (*P. mirabilis*, *S. typhimurium*, *S. aureus*, *L. sakei*, and *C. difficile*), the SiaF protein from *A. timonesis*, and finally the uncharacterised ST5 transporter from *B. thetaiotaomicron* (see Discussion). **(B)** Co-ordination of Neu5Ac by the binding site residues in PmSiaT (adapted from [47]).

**Figure S9.**
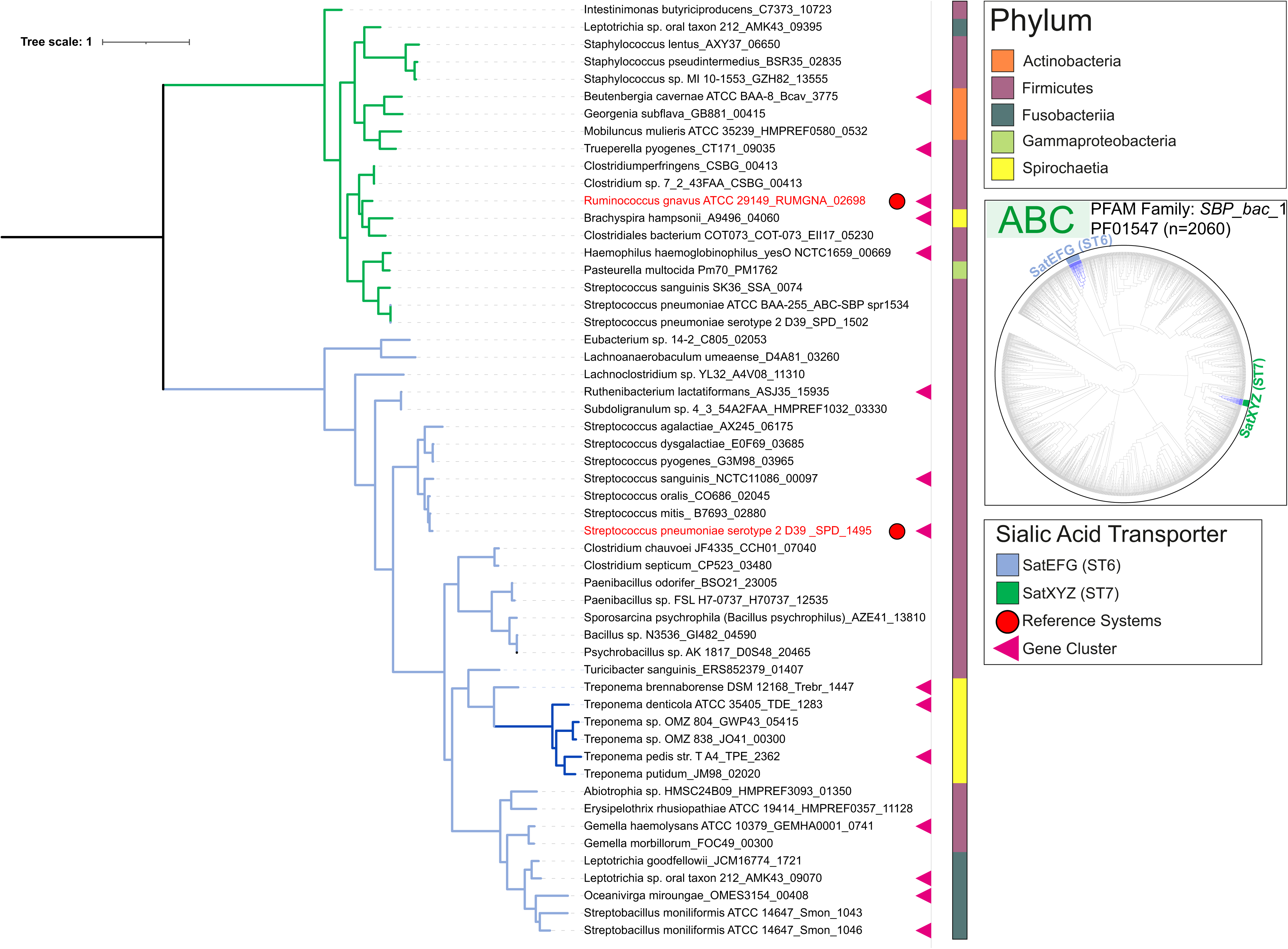
Phylogeny of the ST6-SBP (SatE) and the ST7-SBP (SatX) components of ABC sialic acid transporters. Maximum likelihood phylogeny of sialic acid-binding protein components (SatE and SatX, respectively) of the ST6 and ST7 ABC transport systems. Reference organisms are highlighted in red. *Treponema* spp. possessing “orphan” *satE* genes linked to genes for methyl-accepting proteins (see Text) are marked out by dark blue branches.

**Figure S10.**
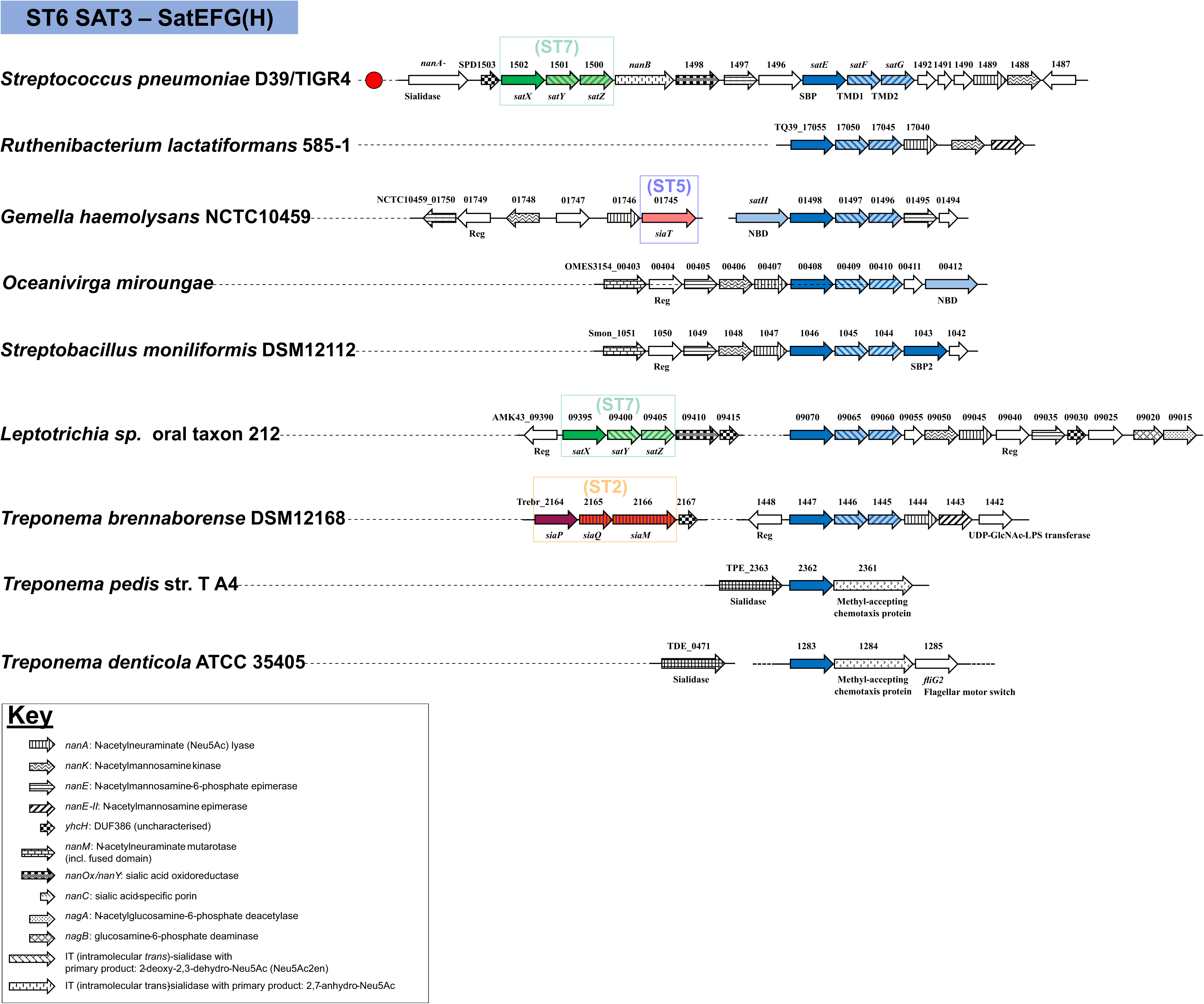
Organisation of ST6 (*satEFG*) gene clusters in selected bacteria. ST6 transporter genes are colour-coded, patterns are used to distinguish among different subunits. Catabolic *nan* genes are pattern-coded and described in the key. For clarity other conserved genes recurring in these clusters are not pattern-coded except when emphasising specific genetic links. The function of other notable genes is reported (without pattern) if relevant for cluster annotation or to exemplify instances of potential HGT. Reference gene clusters are highlighted with a red circle. While ST6 transporter loci normally do not contain an NBD gene, relying instead on multitask *msmK*-like genes for function [44], note here two cases where specific NBD genes have been recruited to the cluster. The incomplete SAT3 cluster of the sialic acid-utilising organism, *Ruthenibacterium lactatiformans* strain 585-1 [67] was completed by matching it to a near-identical locus in the sequenced genome of strain 82B1, which was then used to annotate the remaining genes. Some *Treponema* spp. only contain the *satE* gene coding for the SBP component (Fig. S9), which however possesses a conserved genetic link with a methyl-accepting chemotaxis protein (MCP), suggesting a role in sialic acid sensing and chemotaxis independent of a role in uptake. In *T. denticola*, the ST6 locus also contains an allelic variant of the flagellar motor switch protein gene, *fliG*, consistent with the chemotaxis hypothesis. Reg: regulator gene; SBP: solute-binding protein; TMD: transmembrane domain; NBD: nucleotide-binding domain (ATP-ase protein); HGT: horizontal gene transfer.

**Figure S11.**
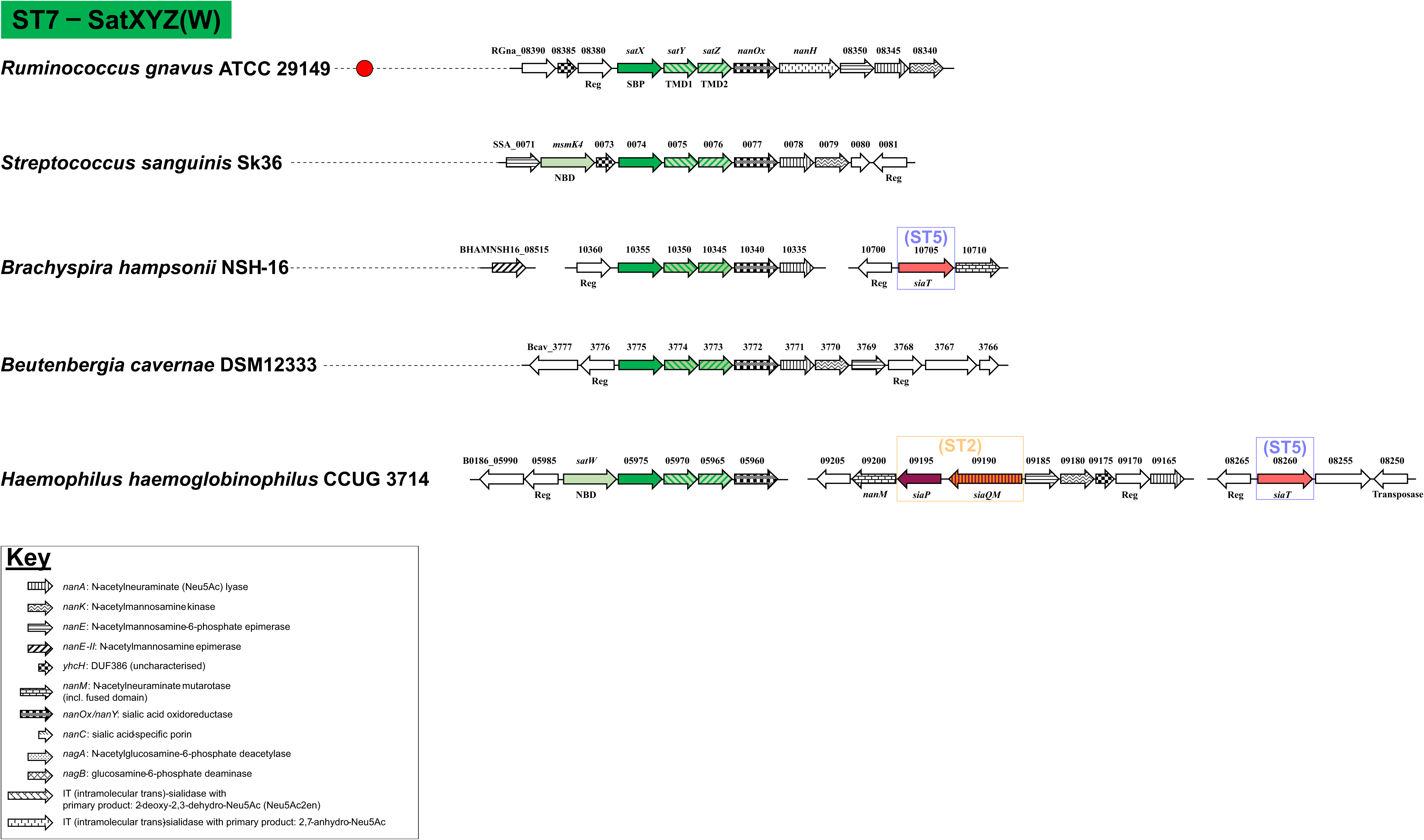
Organisation of ST7 (*satXYZ*) gene clusters in selected bacteria. ST7 transporter genes are colour-coded, patterns are used to distinguish among different subunits. Catabolic *nan* genes are pattern-coded and described in the key. For clarity other conserved genes recurring in these clusters are not pattern-coded except when emphasising specific genetic links. The function of other notable genes is reported (without pattern) if relevant for cluster annotation or to exemplify instances of potential HGT. Reference gene clusters are highlighted with a red circle. While ST7 loci are normally devoid of NBD genes, presumably relying on multitask *msmK*-like for function as shown for *S. pneumoniae* SatEFG [44], note here two cases where different NBD genes have been recruited to the cluster, namely in *H. haemoglobinophilus* (*satW*) and *S. sanguinis SK36* (*msmK4* coding for supernumerary allelic variant of *msmK*). Reg: regulator gene; SBP: solute-binding protein; TMD: transmembrane domain; NBD: nucleotide-binding domain (ATP-ase protein); HGT: horizontal gene transfer.

**Figure S12.**
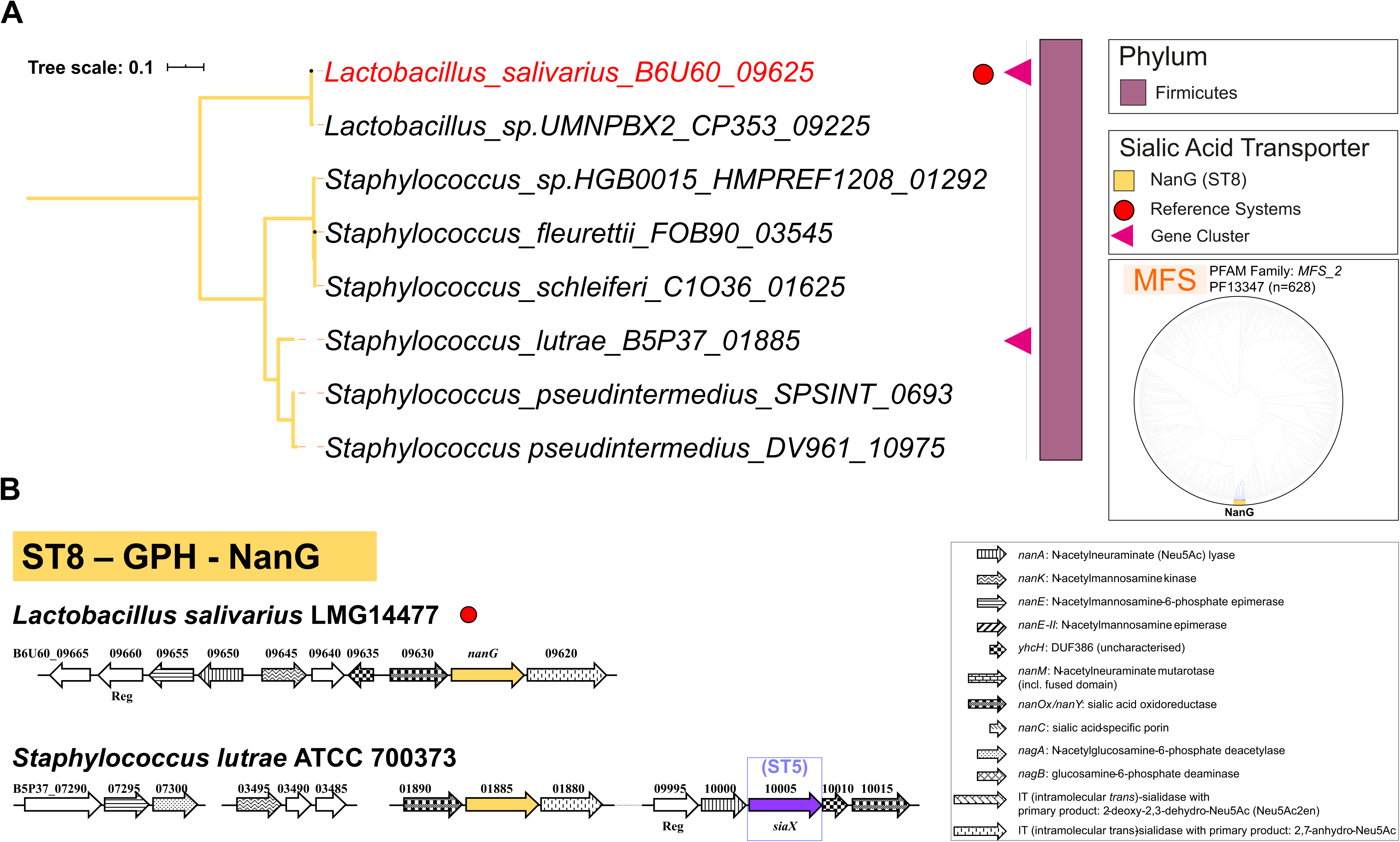
Phylogeny of ST8 (GPH) sialic acid transporters, and organisation of ST8 (*nanG*) clusters in selected bacteria. ST8 transporter genes are colour-coded. Catabolic *nan* genes are pattern-coded and described in the key. For clarity other conserved genes recurring in these clusters are not pattern-coded except when emphasising specific genetic links. The function of other notable genes is reported (without pattern) if relevant for cluster annotation or to exemplify instances of potential HGT. Reference gene clusters are highlighted in red. Note that *S. lutrae* possesses two predicted anhydro-sialic acid transporter genes. Reg: regulator gene; HGT: horizontal gene transfer.

